# Integrating citizen science data with expert surveys increases accuracy and spatial extent of species distribution models

**DOI:** 10.1101/806547

**Authors:** O.J. Robinson, V. Ruiz-Gutierrez, M.D. Reynolds, G.H. Golet, M. Strimas-Mackey, D. Fink

## Abstract

Information on species’ habitat associations and distributions, across a wide range of spatial and temporal scales, are a fundamental source of ecological knowledge. However, collecting biological information at relevant scales if often cost prohibitive, although it is essential for framing the broader context of more focused research and conservation efforts. Citizen-science data has been signaled as an increasingly important source of biological information needed to fill in data gaps needed to make more comprehensive and robust inferences on species distributions. However, there are perceived trade-offs of combining highly structured, scientific survey data with largely unstructured, citizen-science data. As a result, the focus of most methodological advances to combine these sources of information has been on treating these sources as independent. The degree to which each source of information is allowed to directly inform a common underlying process (e.g. species distribution) depends on the perceived quality of the data. In this paper, we explore these trade-offs by applying a simplified approach of filtering citizen-science data to resemble structured survey data, and analyze both sources of data under a common framework. To accomplish this, we explored ways of integrating high-resolution survey data on shorebirds in the northern Central Valley of California with observations in eBird for the entire region that were filtered to improve their quality. The integration of survey data with the filtered citizen-science data in eBird resulted in improved inference and predictive ability, and increased the extent and accuracy of inferences on shorebirds for the Central Valley. The structured surveys were found to improve the overall accuracy of ecological inference based only on citizen-science data, by increasing the representation of data collected from high quality habitats for shorebirds (e.g. rice fields). The practical approach we have shown for data integration can be also be used to improve the efficiency of designing biological surveys in the context of larger, citizen-science monitoring efforts, ultimately reducing the financial and time expenditures typically required of monitoring programs and focused research. The simple processing and filtering method we present can be used to integrate other types of data (e.g. camera traps) with more localized efforts (e.g. research projects), ultimately improving our ecological knowledge on the distribution and habitat associations of species of conservation concern worldwide.

## INTRODUCTION

Information on species’ habitat associations and distributions are a fundamental source of ecological knowledge (Sofaer et al. 2019).This information is often of interest across a broad range of spatial and temporal scales, from high-resolution information that is more relevant for research on habitat selection (Matthiopoulos et al. 2011) or needed to inform management objectives (Zipkin et al. 2010) to larger-scale inferences that are useful to address broader questions (e.g. potential range-shifts with changing climatic conditions; Lyon et al. 2019). However, the process of collecting biological observations across large-spatial scales is often cost-prohibitive for most research and monitoring efforts. Under the best-case scenario, researchers and practitioners are able to monitor plant and animal communities just within their study regions, and during specific times of year. However, the need to make inferences beyond the sampled range of environmental conditions and seasons often limits our understanding of the boarder context of our results, and can limit the use of applied research to inform future monitoring efforts and effective conservation actions.

Citizen-science data has been signaled as a promising source of information to fill-in information gaps needed to model species distributions (Bradter et al. 2018, Gouraguine et al. 2019). However, there are few examples of the potential trade-offs of combining observational data collected at more localized scales with large-scale citizen-science data. This might be due in part to inherent differences that often exist between the two data types. On the one hand, you may have data that is collected at a high spatial resolution using skilled observers, sampling effort is often standardized, and sampling occurs across a habitat gradient that is representative of the region of interest. On the other hand is citizen-science data, which can be collected across a wide range of sampling conditions by observers that vary widely in their level of expertise, collected across a wide range of spatial resolutions, and sampling effort is not standardized. This has created a perceived trade-off between data quality and quantity among data collected from structured, scientific surveys and data collected from larger-scale, volunteer-based monitoring efforts (Figure 1). The assumption being made is that an increase in quantity of citizen-science data comes at a significant cost to quality. Therefore, the logical framework to integrate these two sources of information would be one that would treat them as independent sources of information used to inform a common underlying distribution for a given species (Pacifici et al. 2017).

**Figure 1.**
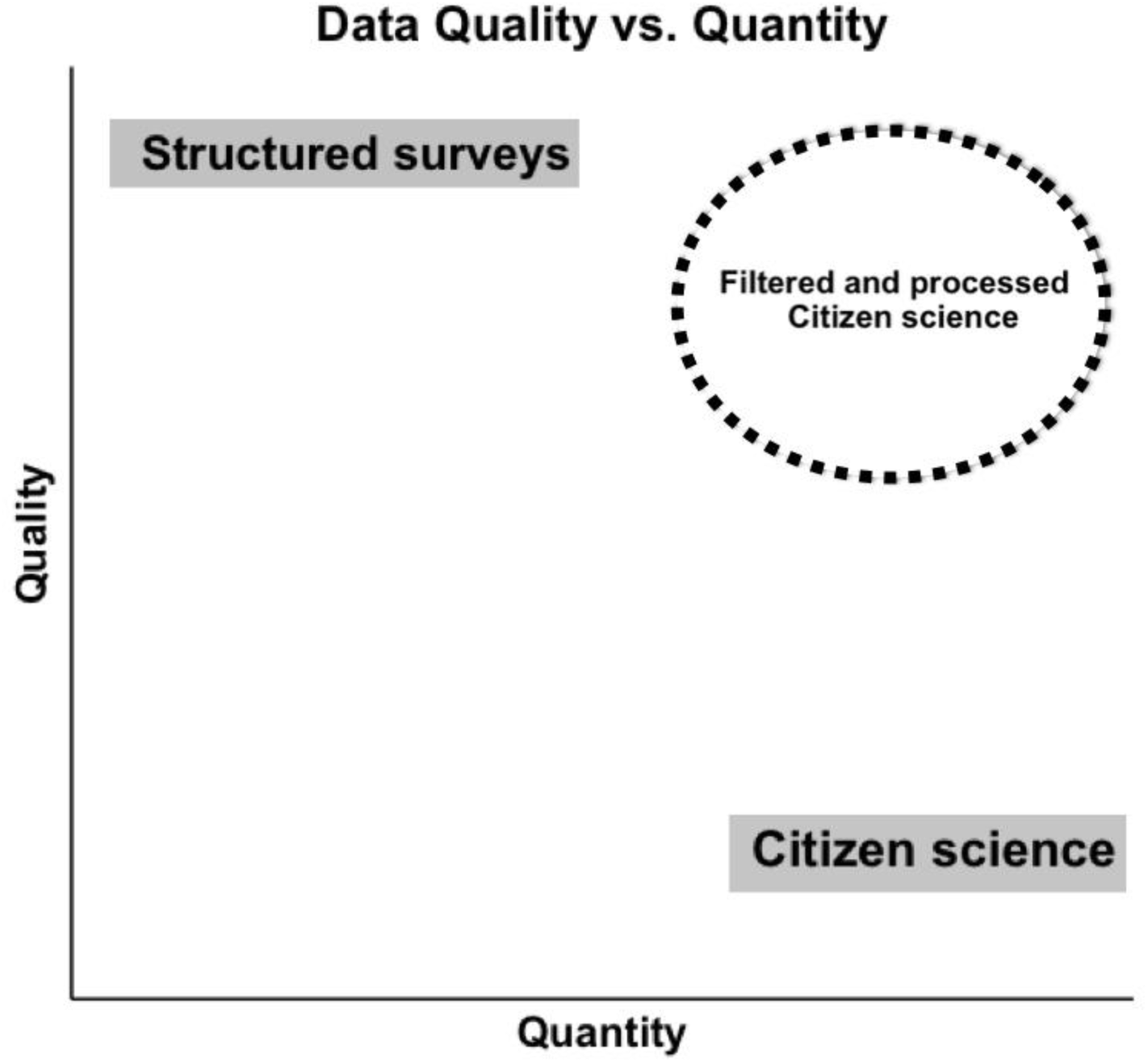
The perceived trade-off between data quality and quantity of data collected using structured surveys and citizen-science efforts. You can bypass this tradeoff by processing and filtering citizen-science data using criteria such as count type (e.g. stationary vs. traveling), duration of counts (e.g. all observations < 30mins), and other measures that align well with the existing data set. This approach is recommended as the most flexible data integration method for the application of a broad-range of species distribution models.

The integration of different data sources is a growing area of methodological development in ecological statistics, and recent advances have been made to develop ways of integrating survey data (e.g. structured) with citizen-science data (e.g. un-structured data) (Miller et al. 2019). For observational data collected at discrete locations, these methods include specifying a joint-likelihood for the two data sources to estimate the underlying species distribution (Miller et al. 2019). In cases where this is not possible, the data source that is deemed as of lower quality (e.g. citizen science data, museum observations) can be used in two ways: 1) modeled as a covariate of the underlying distribution, or 2) used to estimate a separate species distribution, where a correlation structure is specified to share information across data sources. Pacifici et al. (2017) tested these different approaches to integrate observational data from the citizen science project eBird (Sullivan et al, 2017) with more structured data from the North American Breeding Bird Survey (BBS; Sauer et al. 2017). Their results showed that the joint-likelihood approach of combining eBird and BBS data outperformed all other approaches, including using BBS data alone.

The approach used by Pacifici et al (2017) and Miller el at (2019) summarized observational data at a coarse, grid-level in order to account for differences in effort, sampling approach, and other variables that are known to influence detectability (Guillera-Arroita 2017). In addition, Pacifici et al (2017) wanted to reduce potential bias related to the degree of uncertainty about the spatial scale that observations were collected for each independent eBird checklist. This mismatch in scales between the two data sources is what often makes data integration between high-resolution survey data (e.g. point count observations) with lower-resolution citizen-science data non-trivial. However, many citizen-science programs collect high-scale resolution information (e.g. camera traps) in ways that we can infer absences, and collect additional information on effort (e.g. distance traveled, number of hours sampled) that is highly valuable for improving the accuracy of SDMs (Kelling et al. 2019).

Here, we explore a practical approach for data integration between high-quality, citizen-science data with structured survey data that builds upon existing methods for ‘data pooling’ (e.g. Fithian et al. 2015). We explore the trade-offs in inference of using citizen-science data alone, more localized and structured data alone, and pooling together both data sets combined. In addition, we examine specific trade-offs when combining structured and un-structured data sources by exploring the performance of increasing the quantity of citizen-science data through simulations, vs. the addition of data from more targeted survey efforts. To accomplish this, we use The Nature Conservancy’s (TNC) BirdReturns project as a case study (Reynolds et al. 2017). We explore ways of combining high-resolution bird survey data collected for shorebirds on rice fields in the northern region of the Central Valley in California, with observations in eBird for the entire Central Valley that are filtered to improve their quality. The specific aim of the case study is to provide a framework for leveraging survey data with citizen-science data to build more accurate distribution models for shorebird species across the extent of the Central Valley.

## METHODS

### Data

We used point counts carried out during spring surveys (February 1-May 31; n = 8,192) as part of the TNC BirdReturns project conducted in 2014-2017. This project used predicted shorebird occurrence and abundance (Johnston et al., 2015) along with predicted surface water in the Sacramento River Valley to identify times and locations that were likely to be important for migrating shorebirds. TNC used a reverse auction approach to select and incentivize rice farmers in the identified locations to flood their fields during the spring and fall, making temporary wetlands available to the migrating shorebirds. Observers made point counts at fields enrolled in the program and at unenrolled control sites, surveying a semi-circle with a 200 m fixed-radius. Each site was surveyed for at least two minutes and lasted as long as necessary to count all birds present (for more detail on count methods see Golet et al. 2018). Effort for each point count consisted of date, time started and ended, and name of observer.

We combined the point counts with data from the citizen science project eBird (Sullivan et al. 2014) collected during the same time period as the point counts. The eBird data was restricted to the Central Valley of California, USA, and to complete checklists so that non-detection could be inferred (Johnston et al. 2019). We also restricted the eBird data to stationary checklists and traveling checklists limited to 300 m. After filtering this data, we were left with 12,891 checklists. Effort variables for the eBird data set were date, time observations started, duration of observation in minutes, survey protocol (stationary or traveling), distance traveled in meters, and number of observers.

We calculated the effort variables from the point counts to match those of the eBird data set. Observer name in the point count data set was converted into number of observers (however it was always one for this study), time started and ended for the point counts was used to calculate duration in minutes, and each point count was treated as a stationary count (distance traveled = 0). By doing this, the two data sets contained the same effort information and were identical in structure which allowed us to simply join them into one combined data set. We added a variable to the combined data set to note whether an observation was a TNC point count or an eBird checklist. We also calculated a checklist calibration index (CCI) for each checklist in the combined dataset to account for variation in expertise among observers (e.g. expertise score; Johnston et al. 2018). We attached the remotely sensed Cropland Data Layer (CDL; Boryan et al. 2011, Han et al. 2012) to the combined data set by computing the percent of each land cover or crop type in the CDL that was present within a 300 m radius centered on each point count or eBird checklist. We also similarly attached cloud-filled data from Water Tracker (Reiter et al. 2018), a high spatial and temporal resolution surface water tracking system for the Central Valley of California.

A comprehensive data set was created using all of the above eBird checklists (12,891) plus simulated eBird checklists equal to the number of TNC point counts that were included in the combined dataset (8,192). This resulted in a dataset that was the same magnitude as the combined dataset and allowed us to determine if the improvement in accuracy from combining the datasets was simply a function of increasing the sample size. The simulated eBird data was created by slightly jittering the spatial covariates of 8,192 randomly chosen checklists, similar to the oversampling procedure in Fink et al. (2019), however we used all checklists rather than only positive observations here. This ensured that we were maintaining roughly the same prevalence rate in the dataset, and also were not simply making exact copies of the randomly chosen checklists.

### Spatial Filtering and Class Imbalance

As spatial bias is always a concern when using citizen science data (Geldmann et al. 2016), we spatially subsampled the combined data set. The data was sparse for many species (proportion of detections for a species < 0.05 for all checklists in the dataset), so class imbalance was also a concern (He and Garcia 2009). We spatially undersampled the data (e.g. Robinson et al. 2017) by first creating a hexagonal grid of 3.5 km cells over the region from which our observations came via the dggridR package (Barnes et al. 2018) in R (R Core Team 2019). We then split the data for a single species into checklists on which the species was detected (positive observations) and those on which the species was not detected (negative observations). We selected one checklist from the negative observations from within each grid cell and recombined the filtered negative observations with the positive observations. As almost all of the spatial bias was from the negative observations (i.e. only a small percentage of the total number of observations were positive observations), this procedure relieves much of the spatial bias, and because only negative observations were filtered out of the dataset, class imbalance is also addressed here (King and Zeng 2001, Robinson et al. 2017).

To alleviate the effects of class imbalance, after spatially sampling eBird checklists for training distribution and population trend models, Fink et al. (2019) oversampled eBird checklists for species that had a prevalence rate of less than 25%. After spatially undersampling our data for the current study, class imbalance was still a concern, as our undersampling improved the imbalance, but it did not improve class balance to 25% for species other than Yellowlegs. Following the recommendation of Fink et al. (2019), we oversampled the positive observations for each of our species below 25% prevalence before training the distribution models. We used the synthetic minority oversampling technique (SMOTE; Chawla et al. 2002) to create one new example of the positive class in the training dataset for every positive observation in the spatially undersampled dataset. The SMOTE algorithm does not create exact copies of the positive class as in traditional oversampling, but instead creates examples of the positive class that occupy the parameter space between a randomly chosen positive observation and a nearest neighbor. We did not randomly undersample our data as is recommended when using SMOTE (Chawla et al. 2002) because our data had already been spatially undersampled as described above. The spatial undersampling and SMOTE procedure was done to each of the datasets. As the spatial undersampling randomly chooses from negative observations within a grid cell, many negative observations may not be included in training the model. Therefore, we sampled the data using this procedure 100 times, creating 100 unique datasets.

### Analysis

We selected and spatially subsampled (but did not oversample) 15% of the combined dataset to be the testing data for evaluation of each model. This test set was selected because our goal is to make accurate predictions across the entire Central Valley, and give equal importance to each location. Therefore, the test set must include observations across the spatial extent of evaluation and be spatially balanced. For the training datasets, we removed any checklist or point count that was in the test set.

We used the R package ‘ranger’ (Wright et al. 2019) to train a random forest model for each of the seven species (or combined species; Table 1) and for each of four datasets: 1) TNC point counts alone, 2) eBird checklists alone, 3) an oversampled eBird dataset, 4) the combined TNC and eBird dataset. For each species, 1000 trees were grown in the ensemble and the number of variables from which the model could select at each split for each tree was set to the square root of the number of variables included in the model (n = 12 ≈ √142). We evaluated the accuracy of the models using multiple predictive performance metrics (PPMs). We did not evaluate the models using the out of bag (OOB) metric calculated by random forest internally as it favors total accuracy over correctly predicting the minority class, and is highly sensitive to class imbalance. We used the test dataset to evaluate mean squared error (MSE) between the model predictions of presence or absence and the true presence or absence in the test set. We also evaluated error using Brier score (Brier 1950), the mean squared error of the probabilistic model predictions and the true presence or absence in the test set. We evaluated each model’s ability to rank positive observations higher than negative ones using the area under the curve (AUC; Fielding and Bell 1997). We evaluated each model’s ability to predict presence or absence using Cohen’s Kappa (Kappa; Cohen 1960) and its components, sensitivity (true positive rate), and specificity (true negative rate). We produced distribution maps for each species and dataset, and we recorded the predictor importance metrics from models trained on each dataset.

**Table 1.** List of species for which we modeled spring distribution in the Central Valley of California with each of the four datasets.

| Common Name | Latin Name | Abbreviation in Tables & Figures |
| --- | --- | --- |
| American Avocet | *Recurvirostra americana* | AMAV |
| Dunlin | *Calidris alpina* | DUNL |
| Greater Yellowlegs* | *Tringa melanoleuca* | YLEG* |
| Least Sandpiper | *Calidris minutilla* | LESA |
| Lesser Yellowlegs* | *Tringa flavipes* | YLEG* |
| Long-billed Curlew | *Numenius americanus* | LBCU |
| Long-Billed Dowitcher** | *Limnodromus scolopaceus* | DOWI** |
| Short-billed Dowitcher** | *Limnodromus griseus* | DOWI** |
| Western Sandpiper | *Calidris mauri* | WESA |

\* Combined in analysis and referred to collectively as “Yellowlegs” in this study
\*\* Combined in analysis and referred to collectively as “Dowitcher” in this study

## RESULTS

The fields that are stored as part of the eBird project allowed us to filter the data to be of a similar protocol as the TNC point counts. This filtering reduced uncertainty in the location of eBird checklists and eliminated the need for coarse level summaries of the data for integration (Pacifici et al. 2017, Miller et al. 2019). For all species, the combined dataset had higher predictive accuracy than either the TNC point counts or eBird checklist datasets on their own; however error (MSE, Brier score) was relatively low and AUC was relatively high for all species and all datasets, particularly for the three datasets where eBird checklists were included (Figure 1; S1). Improvement in accuracy varied among species, however; for the Kappa statistic (predicting presence and absence against the test set), the combined dataset was an improvement over all three of the other datasets evaluated (4.5% - 35.5% improvement; Figure 2). The improvement or loss in the error statistics was usually negligible (with the exception of LBCU; Figure 4; S1) for the combined dataset versus the next best dataset; however, error metrics were already very low for most species, so great improvement here was not likely. Likewise, AUC was relatively high for most species and for the three datasets containing eBird checklists, therefore the gain was negligible for many species (improvement of −0.60% - 7.5%; S1). For the few species/metrics combinations where the combined dataset was not the best performing model, it was the second best, with the best being the eBird checklists with simulated eBird checklists added. The TNC point counts, nor eBird checklists alone performed better for any metric than the dataset where the two were combined.

**Figure 2.**
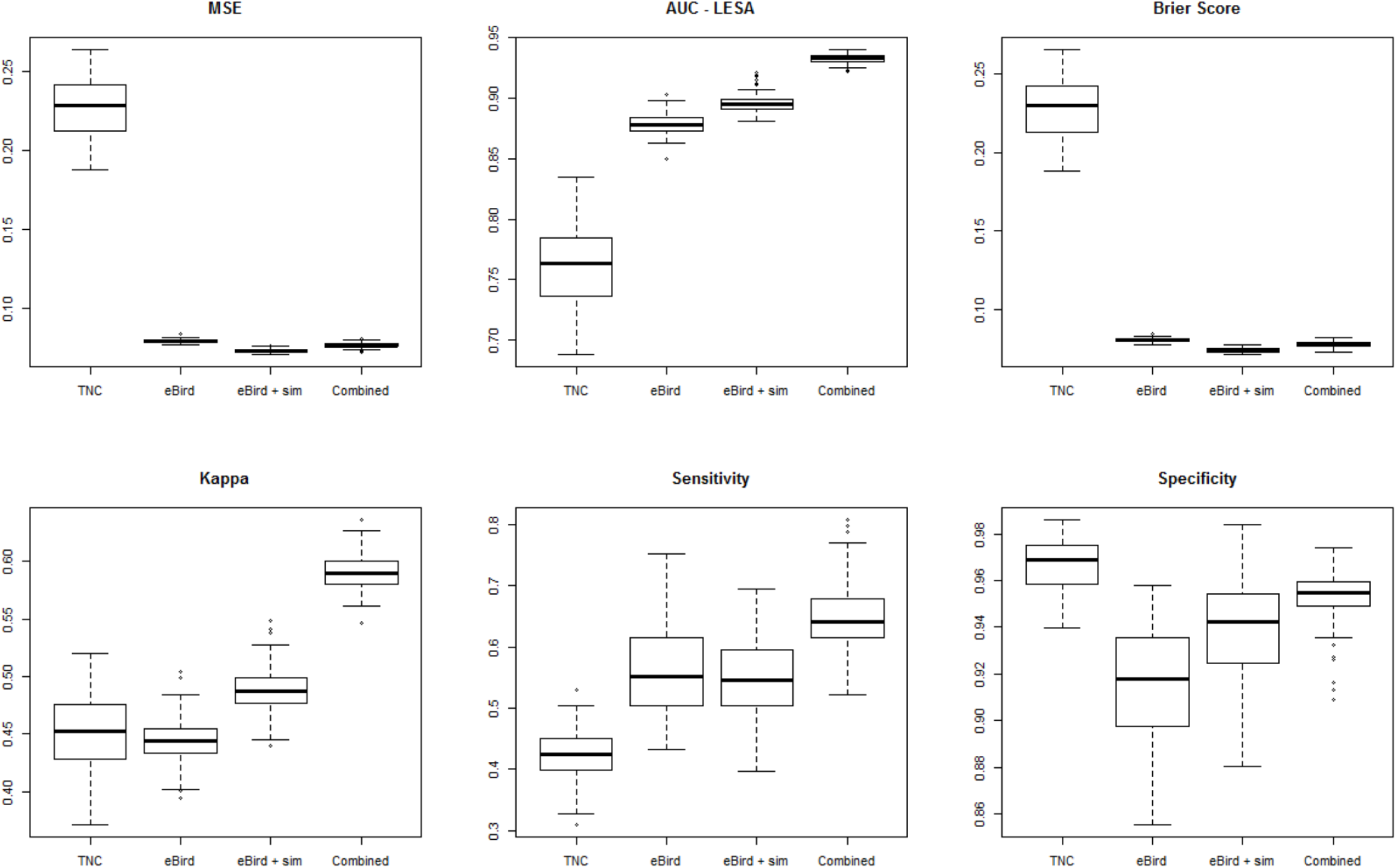
Boxplots for MSE, AUC, Brier score, Cohen’s Kappa, Sensitivity and Specificity for models run for Least Sandpiper (LESA) with each dataset; TNC point counts (TNC), eBird checklists (eBird), eBird checklists with additional simulated eBird data (eBird + sim), and the dataset of TNC point counts and eBird checklists combined (Combined).

**Figure 3.**
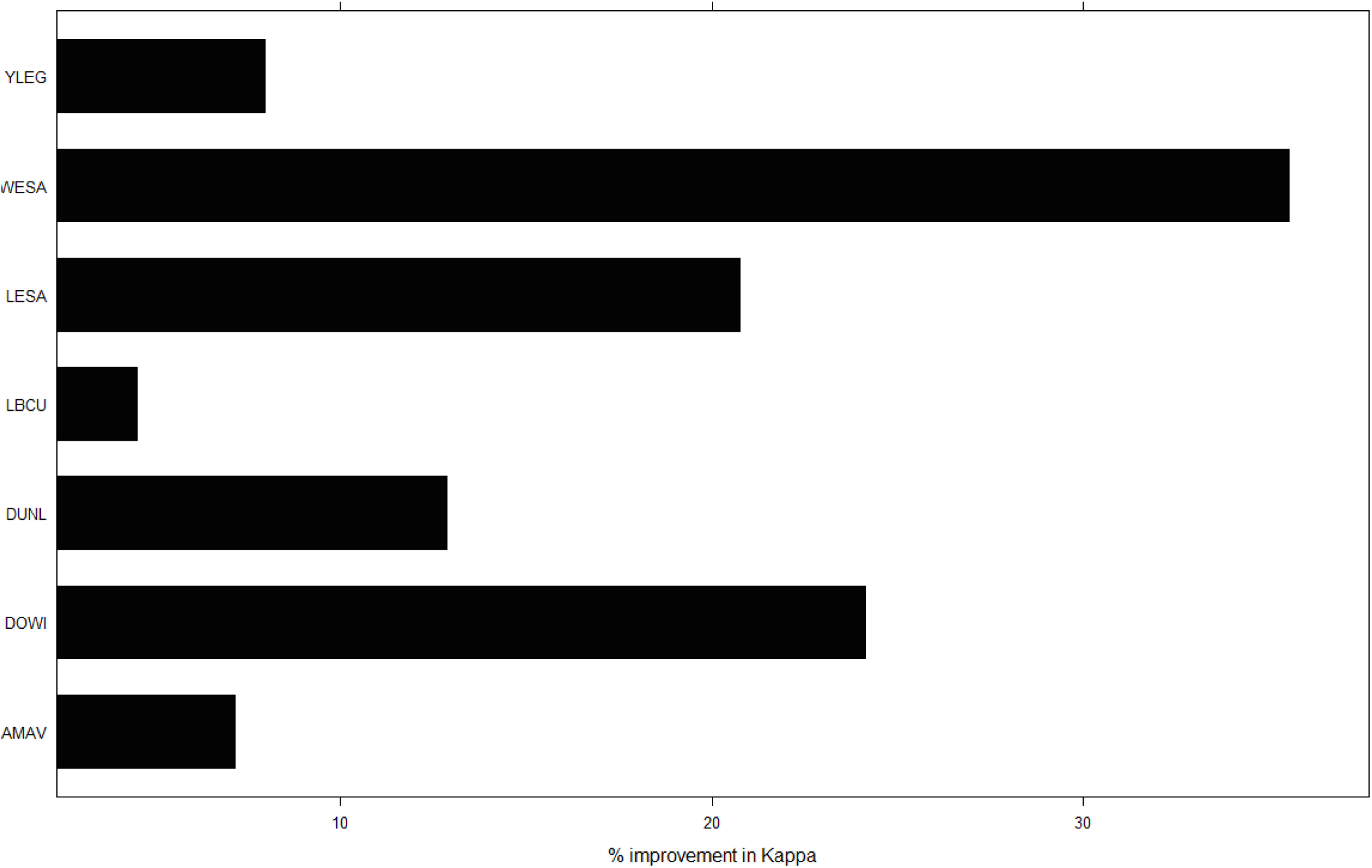
Average improvement in Cohen’s Kappa for distribution models of each species when using the combined dataset versus the dataset that provided the next highest Kappa value.

**Figure 4.**
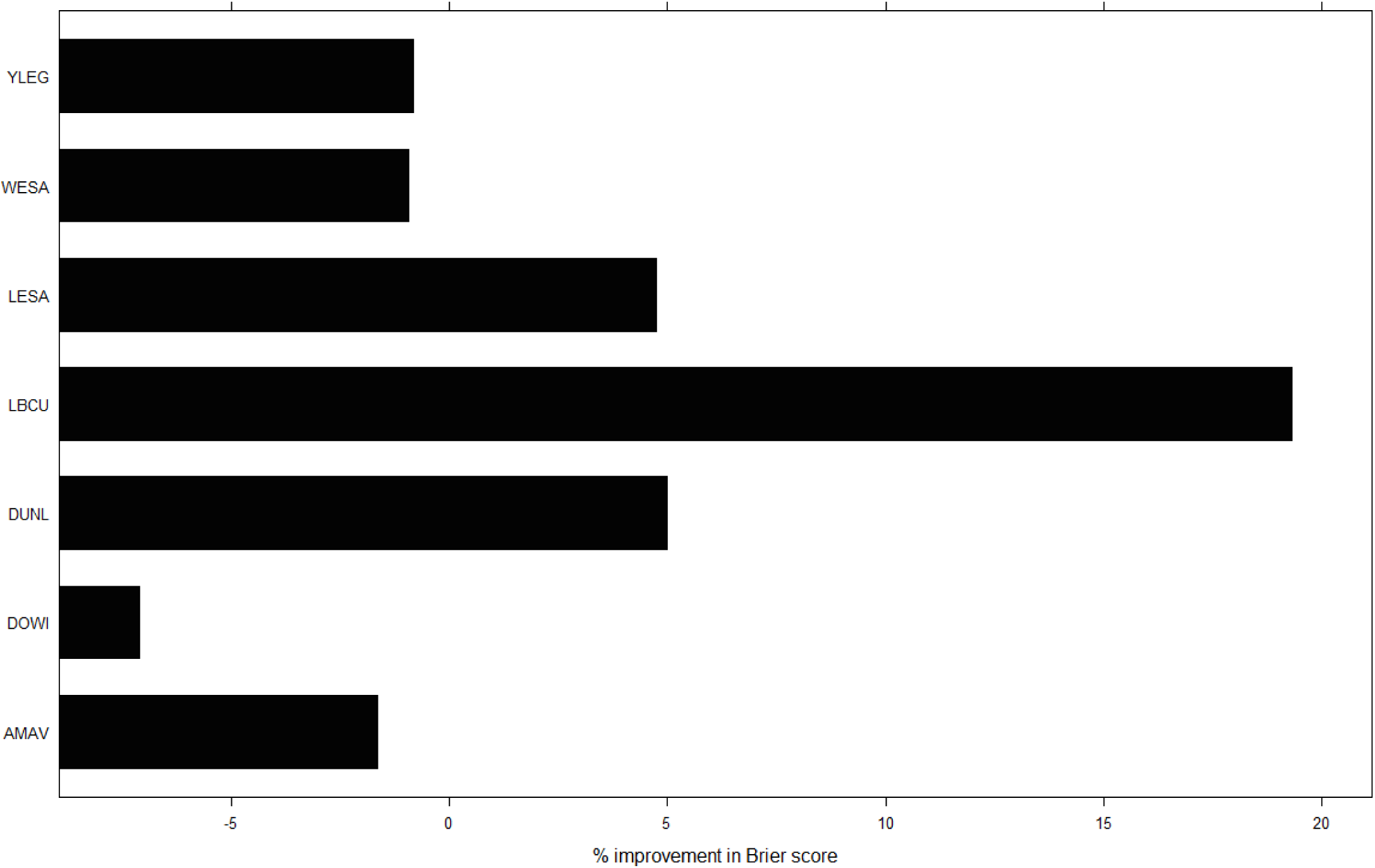
Average improvement or loss in Brier score for distribution models of each species when using the combined dataset versus the dataset that provided the next lowest value.

We produced distribution maps for each species (Figure 5; S1) to determine if there was a difference in the overall pattern of distribution estimated by models trained on the different datasets. For most species, there was greater contrast between the presence estimates and absence estimates for the combined dataset when compared to the other three. Visually, this means hotter ‘hotspots’ and darker regions where absence is predicted (Figure 5; S1). We also collected the important variables for the model when run with each of the datasets. For all species, the importance of rice and water was apparent as it was among the top variables for each of the datasets, however, it was not until the datasets were combined that each rice and the Water Tracker layer had a high importance score (Figure 6; S1). The combination of the data allowed the models to hone in on rice and surface water as highly important, where they had only moderate importance comparatively when using the other datasets. This is likely because only 1.5 % of eBird checklists in our study come from locations where the percent land-cover is >50% rice. Conversely, almost 60% of the TNC point counts come from locations where the percent land-cover is >50% rice.

**Figure 5.**
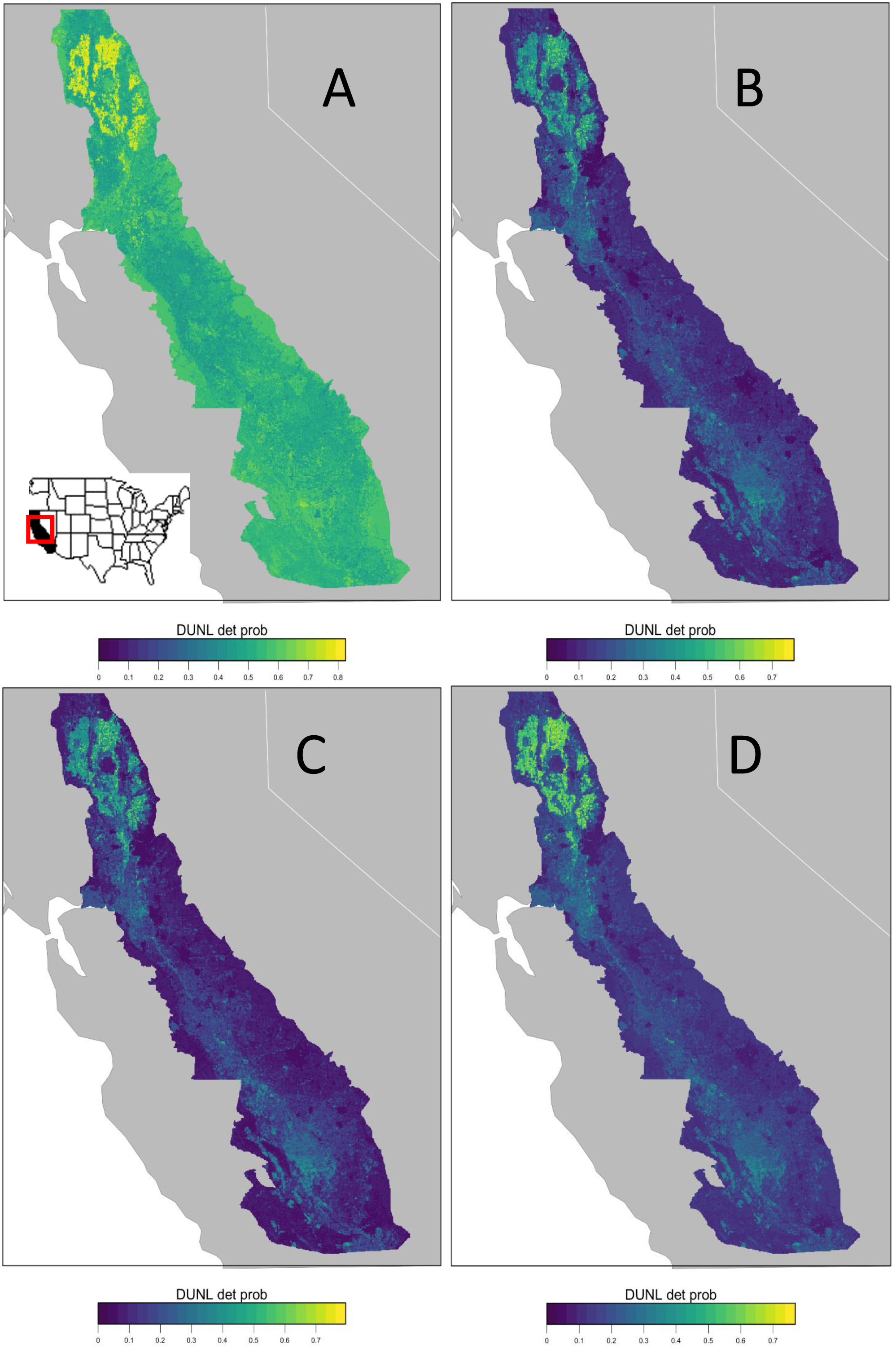
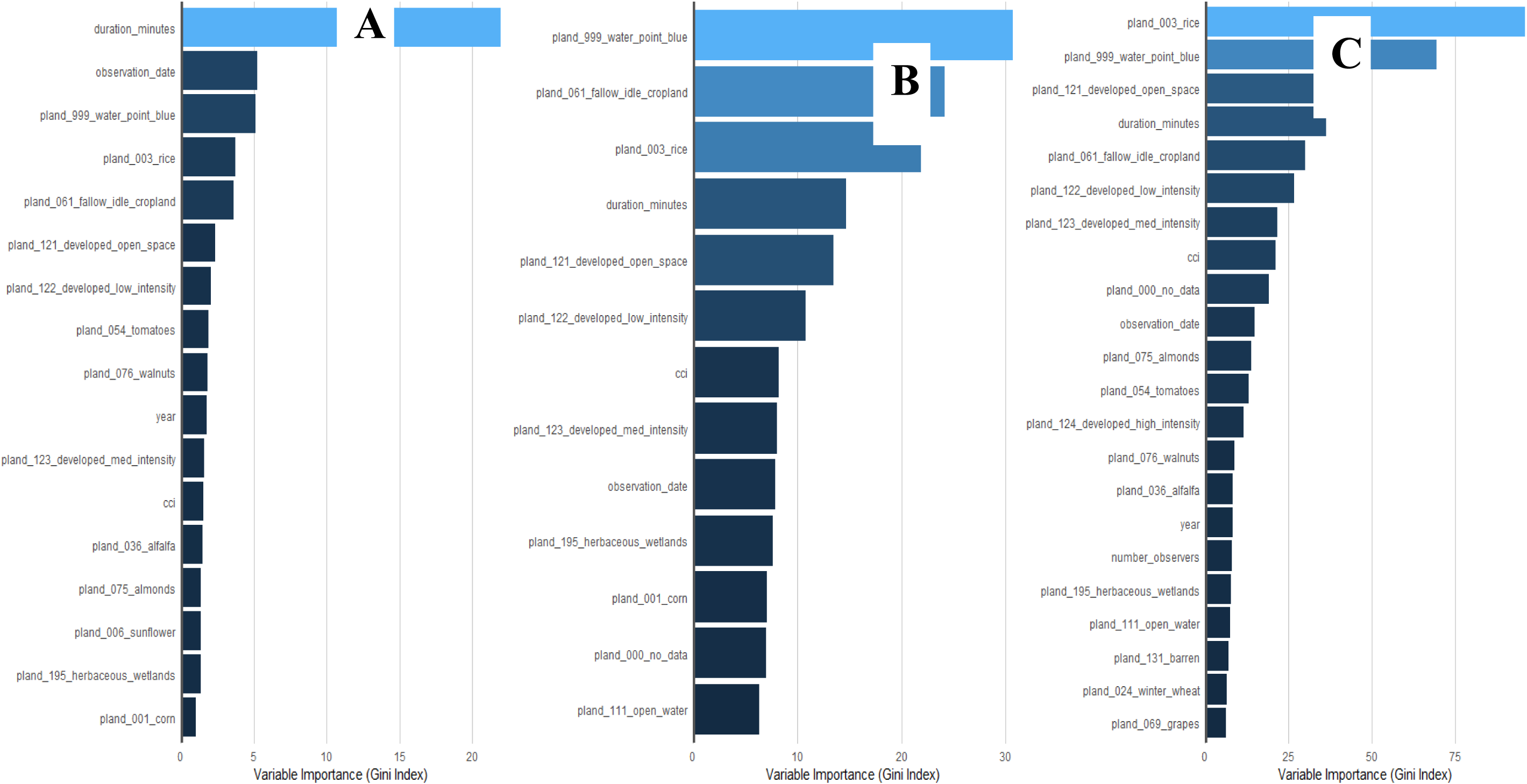
Predicted probability of an expert surveyor detecting a Dunlin in the spring of 2016 on a one-hour long checklist where the distance traveled was < 300m., as estimated by the model using TNC point counts alone (A), eBird checklists alone (B), eBird checklists with added simulated eBird checklists (C), and the combined dataset of TNC point counts and eBird checklists.

## DISCUSSION

Our results lend further support to efforts looking to combine data from multiple sources to improve the inference and/or predictive ability of distribution models (Pacifici et al. 2017, Miller et al. 2019). We have shown that citizen-science data can be filtered to generate a high-quality data set that can closely match the resolution and sampling approach of structured surveys, supporting the call for current and future citizen-science projects to collect essential information related to location and effort, as well as complete surveys (Kelling et al, 2019). The integration of survey data with the filtered citizen-science data in eBird resulted in improved inference, predictive ability, and ultimately increased the extent of inference of the structured surveys. In turn, the structured surveys were able to improve the ecological inference of the citizen-science data, by improving the representation of sampled habitats that are key for shorebird species. Most importantly, the practical approach we have shown for data integration is an improvement on simpler ‘data pooling’ approaches for data integration, and can be used to improve the efficiency of designing biological surveys to collect distribution information in the context of larger, citizen-science monitoring efforts, ultimately reducing the financial and time expenditures typically required of monitoring programs and focused research (Reich et al. 2018).

The combined dataset resulted in improved accuracy across all metrics relative to the TNC survey data or eBird data alone, for all of the species considered in this study. Our combined approach also predicted presence/absence via agreement with a test data set more accurately than the different permutations of datasets considered (e.g. TNC alone, eBird alone, eBird plus simulated eBird). The observed improvement in accuracy is not likely a function of simply increasing sample size, since augmenting eBird data with simulated data did not show the same levels of improvement. The addition of the TNC survey data to eBird data improved the coverage of the data both spatially and in targeted habitats. Previous work has shown that migrating shorebirds heavily use flooded rice fields in the Central Valley, including unflooded rice fields (Elphick and Oring 1998, Golet et al. 2018). Given that rice fields are the main focal habitat of the TNC BirdReturns program, ∼60% of the survey data from this project was carried out at sites where at least 50% of the landcover is rice. In contrast, the majority of the rice fields in the Central Valley are not accessible to regular eBird participants, as they are often on private lands. This is likely why we see so few (∼1.5%) eBird checklists from the Central Valley in locations where at least 50% of the landcover is rice. The addition of more data from high quality habitat is what provided the improvement in accuracy of the combined eBird and TNC data set. This highlights the importance and value of more targeted research and survey efforts within the context of large-scale citizen-science monitoring efforts. Given that private lands make up more than half of the land in the United States, supporting wildlife monitoring efforts on privately held lands that are linked into large-scale efforts such as eBird can greatly improve inferences on species distributions and habitat associations across scales of interest for all stakeholders (Hilty and Merenlender 2003).

Interestingly, models based only on the TNC point counts struggled to learn where each species was likely to be absent across the entire Central Valley because the data came largely from “good” shorebird habitat in the northern portion of the Central Valley. The original intent of this project was not focused on species distribution modeling, so this does not come as a surprise. However, given that absence information allows for more accurate distribution models (Brotons et al. 2004), the addition of eBird data was able to provide information on where species are likely to be absent, and improved inferences on what habitat types most benefit shorebirds, but were not surveyed as part of the TNC monitoring efforts. On the other hand, the models using the eBird data alone were able to predict absences well, and had relatively high accuracy when predicting presence/absence, but overall ecological inference was improved when combined with the point count dataset. The TNC point counts acted as targeted surveys in under-surveyed habitat, which previous work has shown can improve the accuracy of distribution models using eBird checklists (e.g. Xue et al. 2016). The complementary nature of the datasets is also shown when examining the important predictors. For Dunlin (Figure 6), the rice landcover and the Water Tracker layer are important predictors for each dataset, however their importance values more than double when the combined dataset is used to train the model. While other species do not show such a drastic shift in the importance value for these two habitat variables, the pattern is similar for most in that these variables become more important to the model’s predictions when the datasets are combined.

The approach we present for combining data is applicable for other conservation monitoring programs and ongoing research efforts, given that a significant amount of research is conducted over small spatial and temporal scales (Heidorn 2008), and often difficult to scale-up without similarly structured data (Poisot et al. 2019). The data fields that exist in eBird data facilitated further processing and filtering to match the structure of the individual dataset of interest, and allowed us to leverage the strengths of both data sources. However, even citizen-science datasets that do collect effort information are often lacking information on sampling locations, although incentivizing participants to collect data in these data-poor locations has been shown to improve the accuracy of distribution models (e.g. avicaching; Xue et al. 2016). Similarly, data from smaller-scale, individual research projects can also help fill in these gaps in citizen-science data. Given that inferences from small-scale studies cannot be extrapolated to larger spatial extents (Sandel and Smith 2009), the approach we present here for combining data from small-scale studies with citizen-science data filtered to match the existing data structure will increase the overall extent of inference, and improve our ability to conceptualize conservation actions within the larger context of the target population(s) of interest.

Information on species distributions across large scales is one of the most fundamental information needs for basic and applied research fields in ecology. However, this level of information often requires large-scale, coordinated surveys that can be time consuming and costly to manage. In addition, models used to estimate species distributions are often data-hungry, and are often unable to generate information at the spatial and temporal scales that are most relevant for research and conservation efforts. We have shown the utility of combining survey data with semi-structured citizen science data (Kelling et al, 2019) for improving accuracy in species distribution models, which can result in more efficient and cost effective surveys (Pacifici et al. 2017, Reich et al. 2018, Miller et al. 2019). The simple processing and filtering method we present for citizen science data allows for the integration of a small-scale point count data set with eBird checklists, can be used to integrate similar types of data being collected by citizen-scientists (e.g. camera traps) with more localized efforts (e.g. patrolling by park rangers), ultimately improving our ecological knowledge on the distribution and habitat associations of species of conservation concern worldwide.

## Acknowledgements

We would like to thank Point Blue for supplying Central Valley water data. We would also like to thank Tom Auer, Steve Kelling, and Matt Reiter for comments that improved this project and the manuscript.

## Supplementary Information 1

Boxplots for MSE, AUC, Brier score, Cohen’s Kappa, Sensitivity and Specificity for models run for American avocet (AMAV), long- and short-billed dowitcher combined (DOWI), dunlin (DUNL), long-billed curlew (LBCU), western sandpiper (WESA) and greater and lesser yellowlegs combined (YLEG) with each dataset; TNC point counts (TNC), eBird checklists (eBird), eBird checklists with additional simulated eBird data (eBird + sim), and the dataset of TNC point counts and eBird checklists combined (Combined). Accuracy metrics for least sandpiper (LESA) are in the main text.

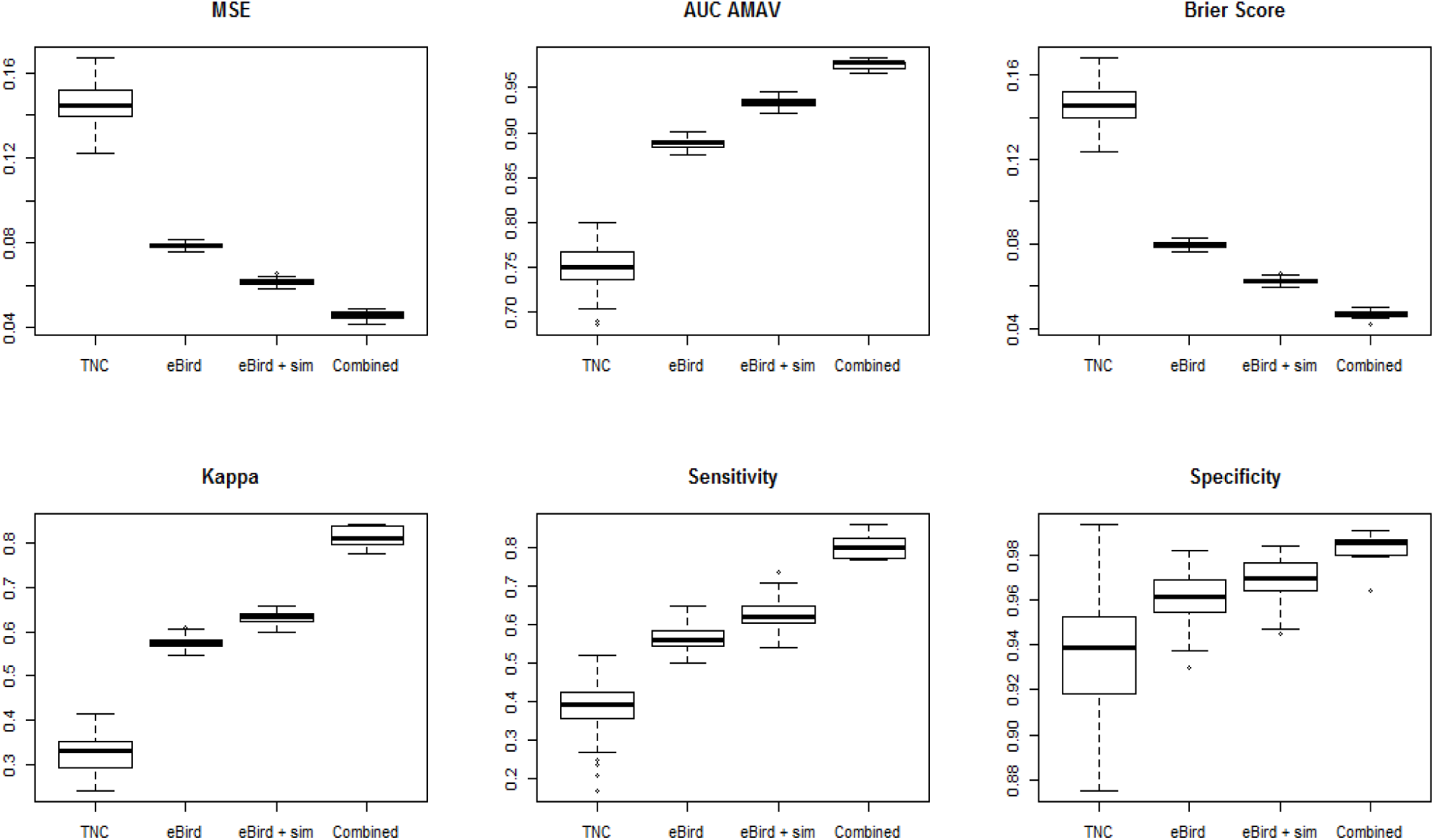

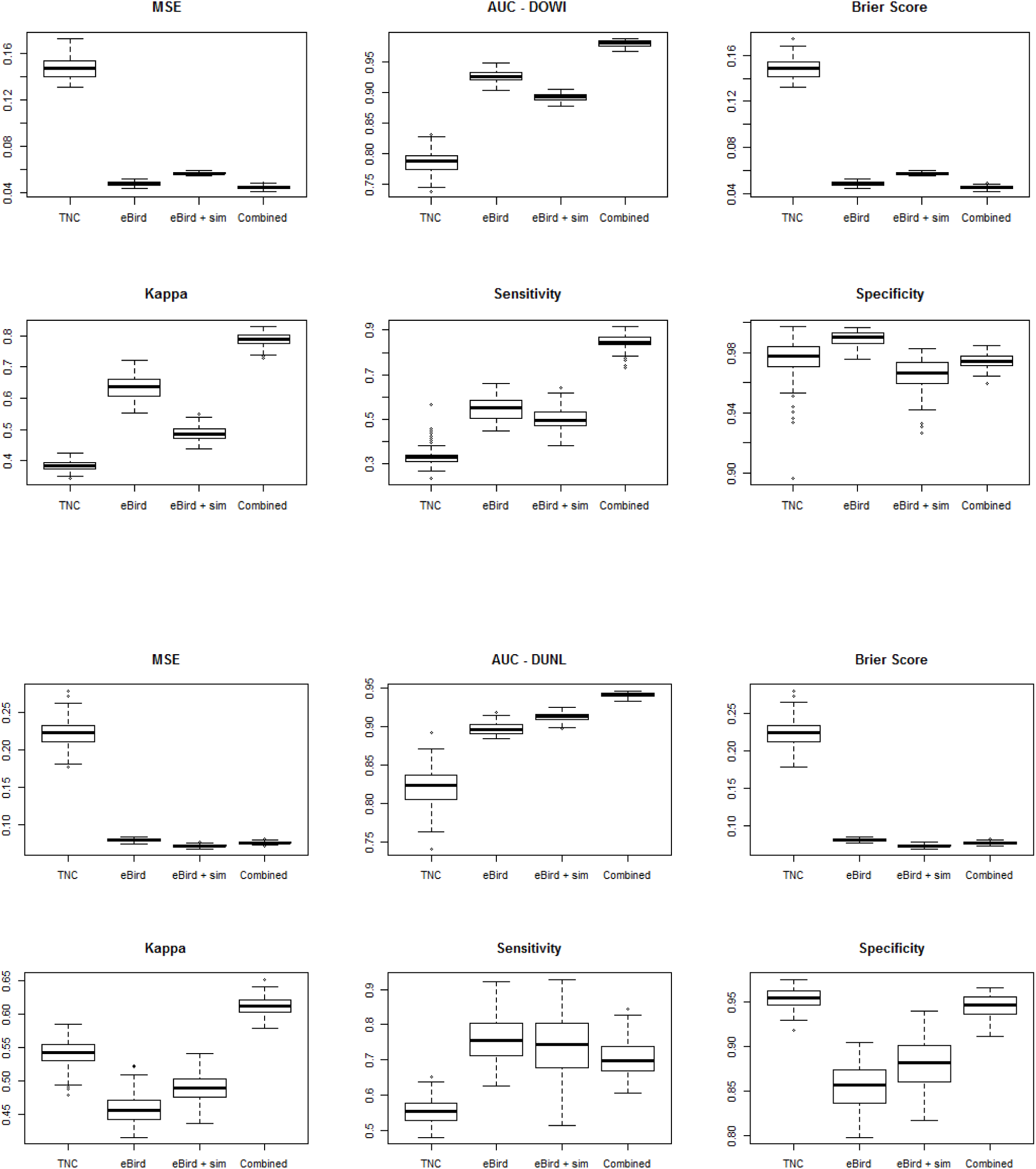

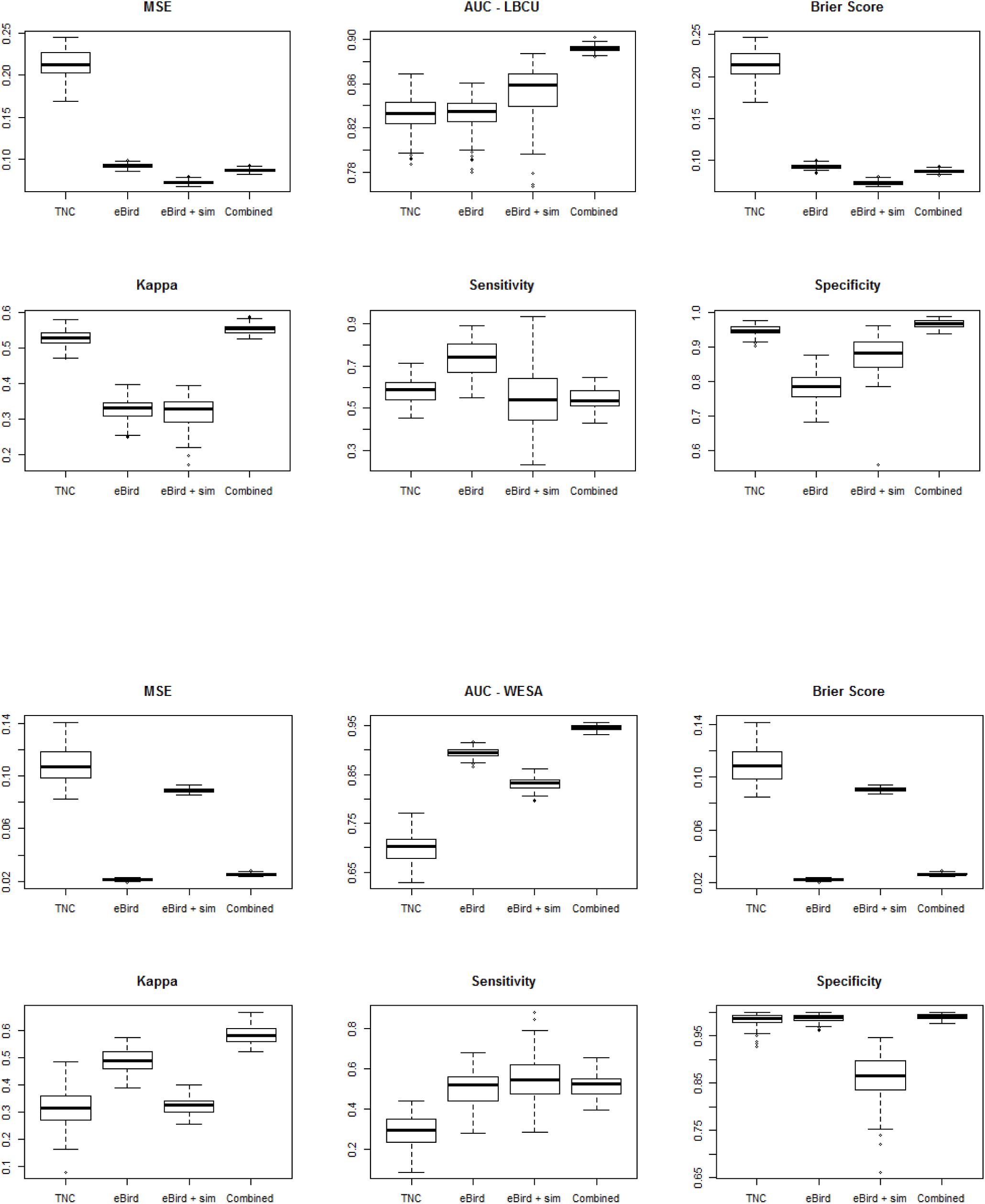

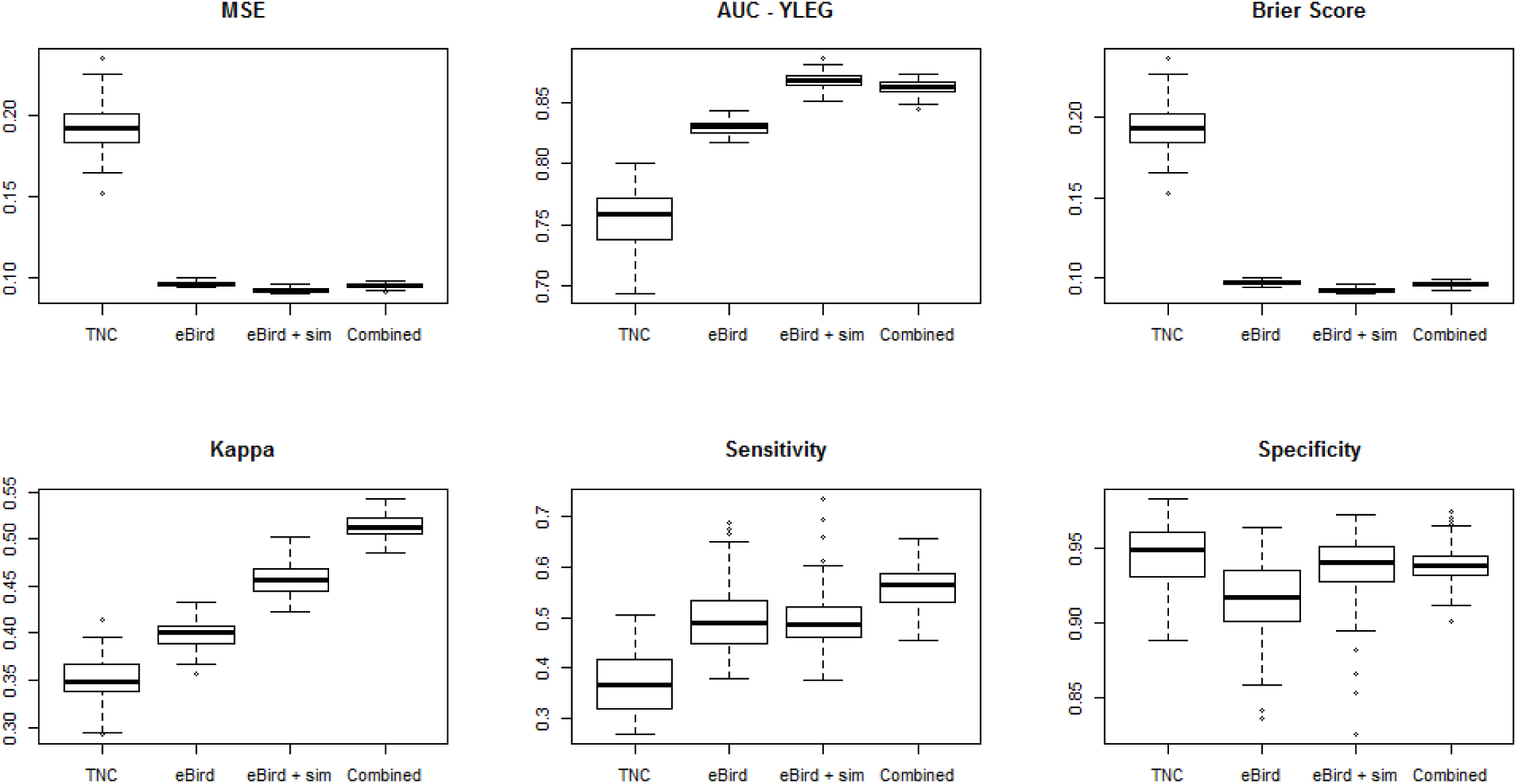

## Supplementary Information 2

Average improvement or loss in mean squared error and AUC for distribution models of each species when using the combined dataset versus the dataset that provided the next lowest value. Recall that the values for these metrics were already very low (for MSE) and very high (for AUC) to begin with. So even a somewhat large change in the percent improvement or loss represents a rather small change for most species. The plots showing improvement or loss for Brier score and Kappa are in the main text.

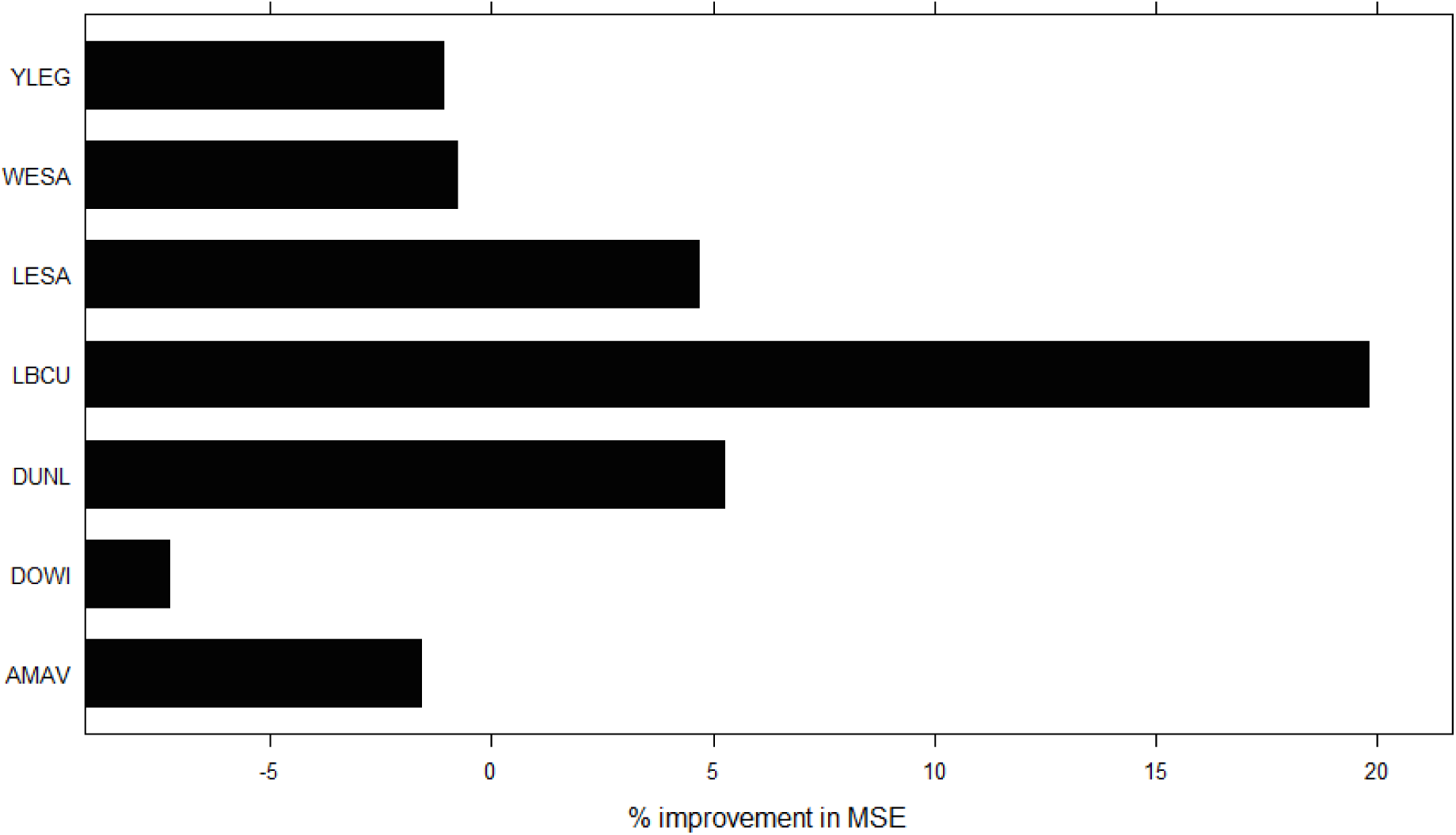

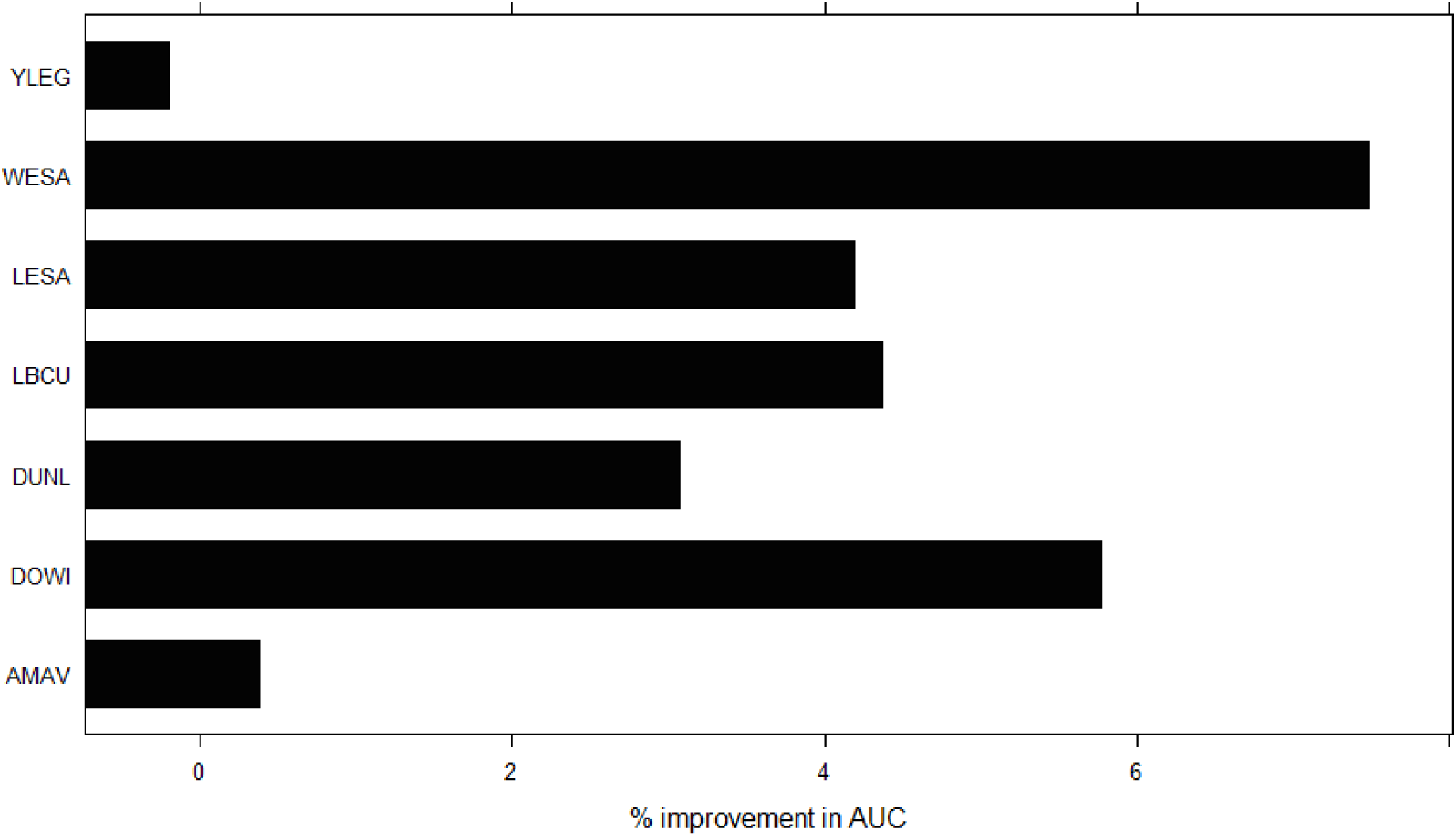

## Supplementary Information 3

Predicted probability of an expert surveyor detecting an American avocet (AMAV), long- and short-billed dowitcher combined (DOWI), dunlin (DUNL), long-billed curlew (LBCU), least sandpiper (LESA), western sandpiper (WESA) and greater and lesser yellowlegs combined (YLEG) in the spring of 2016 on a one-hour long checklist where the distance traveled was < 300m., as estimated by the model using TNC point counts alone (A), eBird checklists alone (B), eBird checklists with added simulated eBird checklists (C), and the combined dataset of TNC point counts and eBird checklists (D). Maps for dunlin (DUNL) are in the main text.

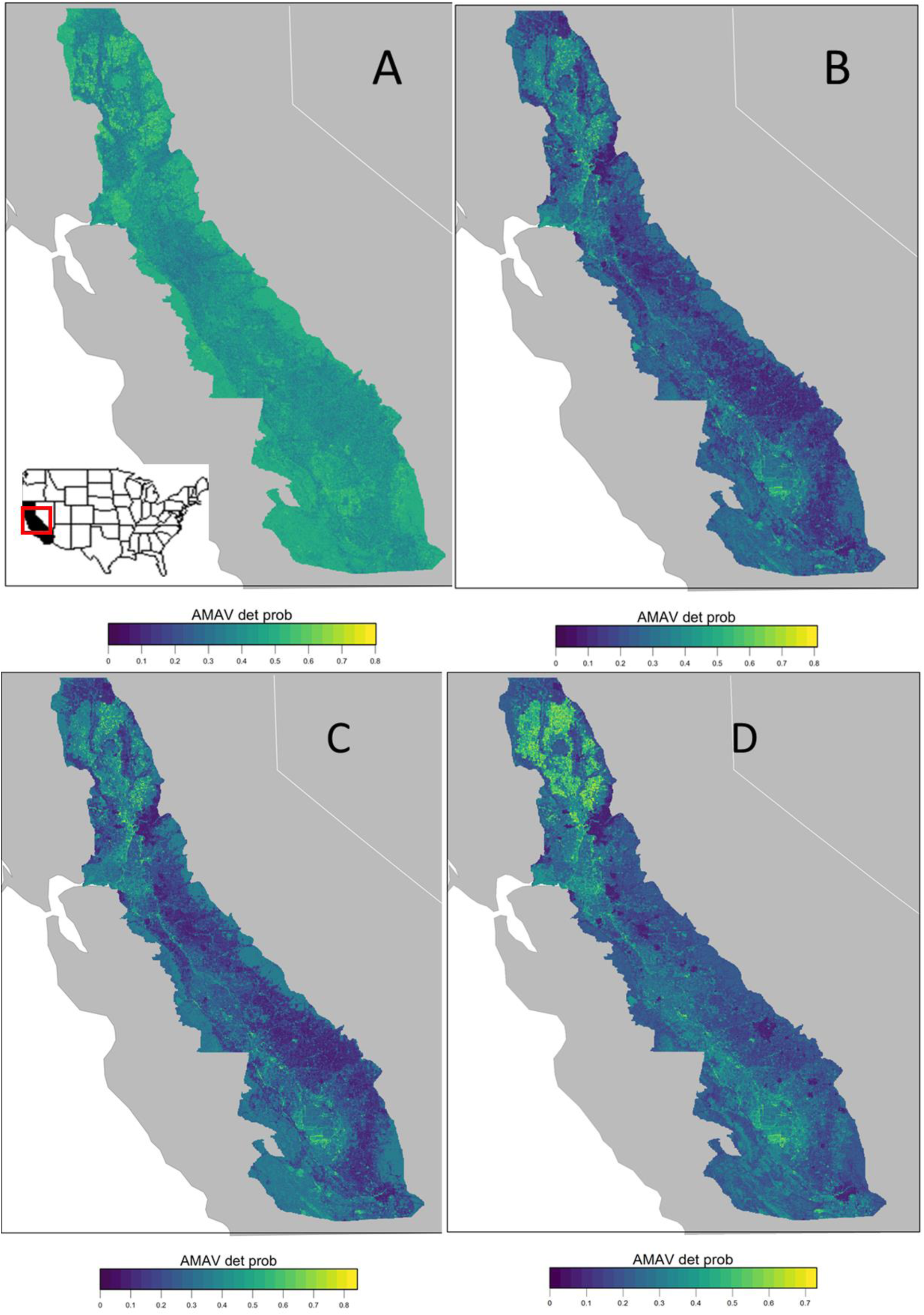

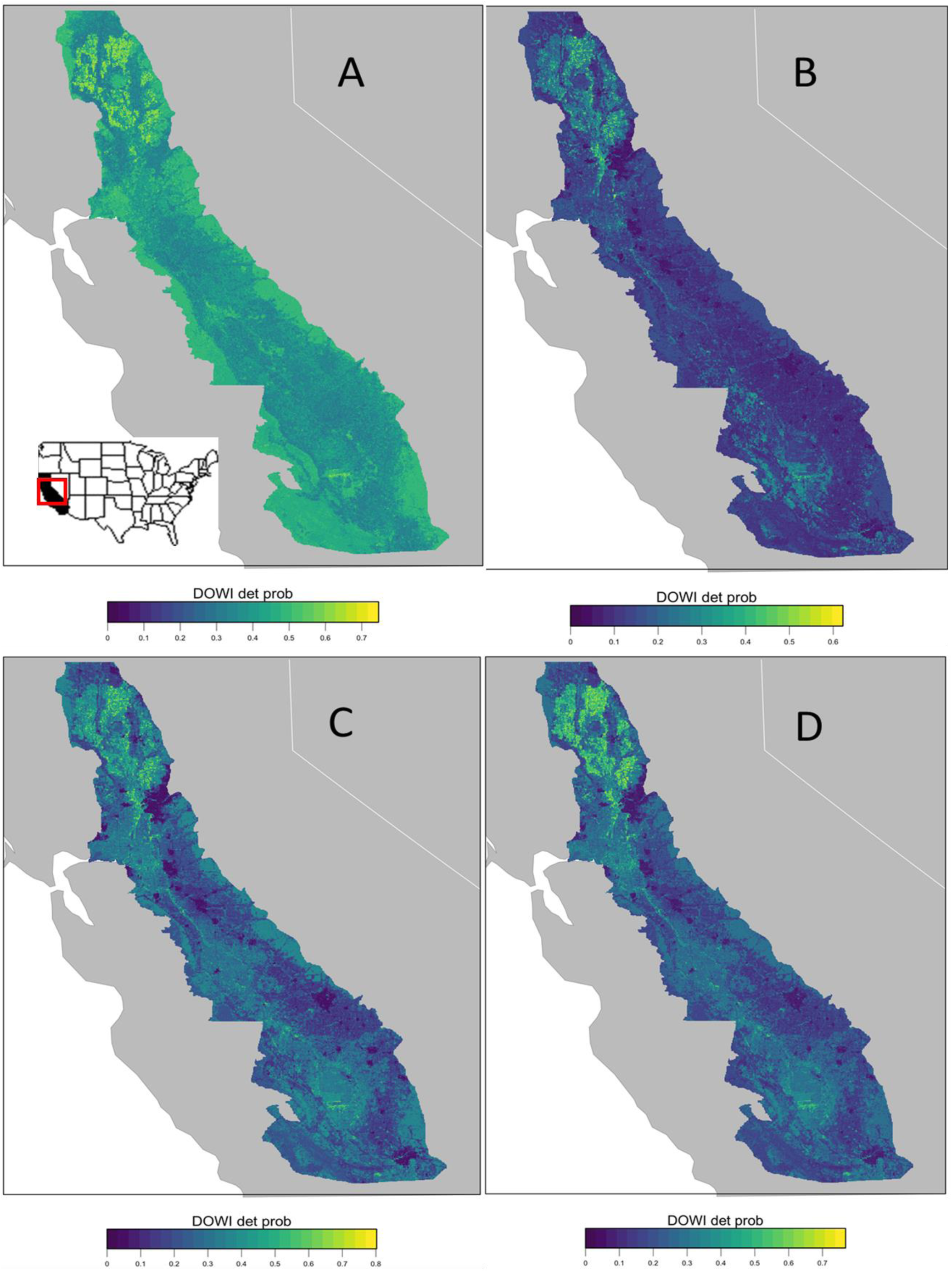

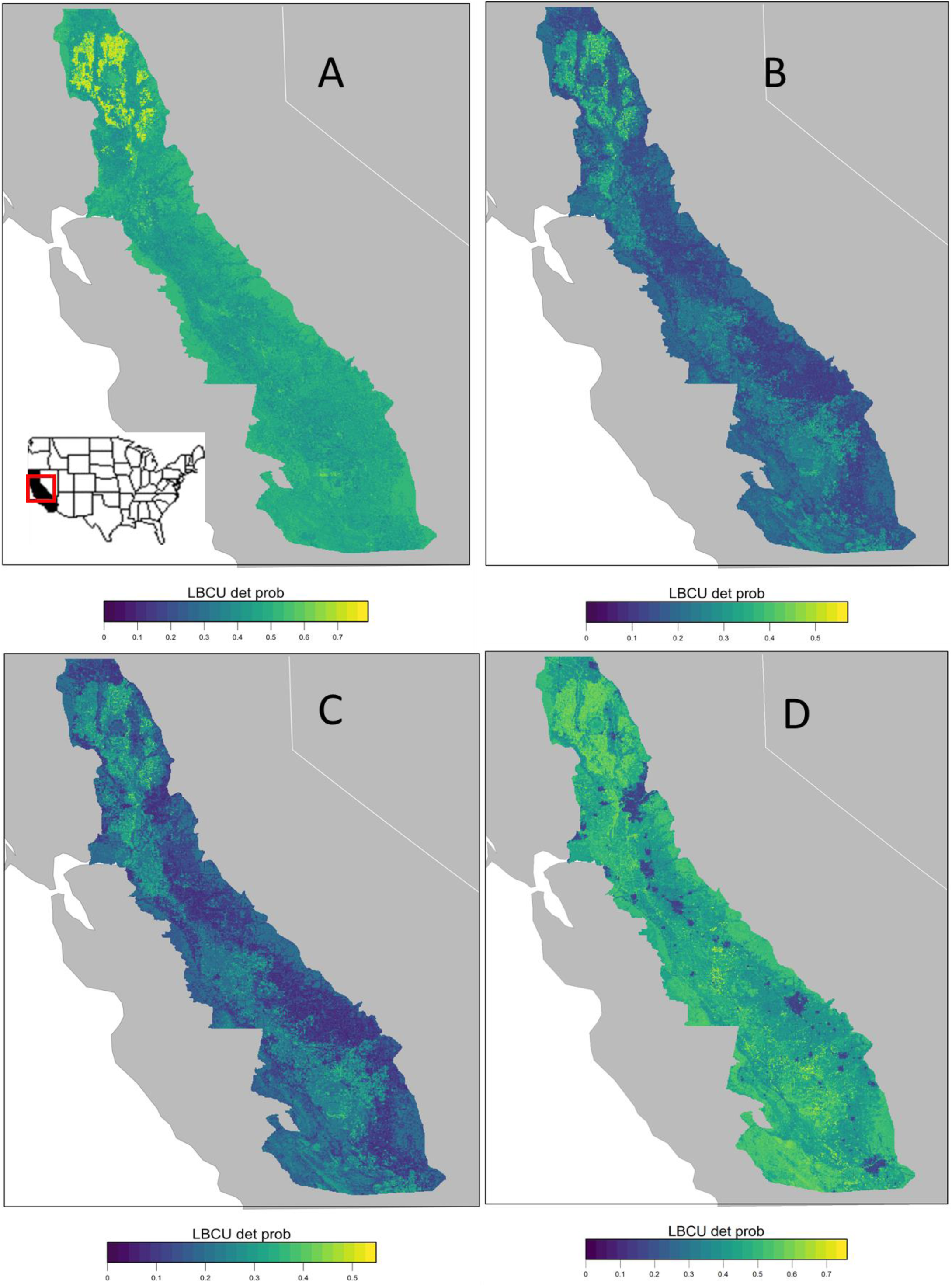

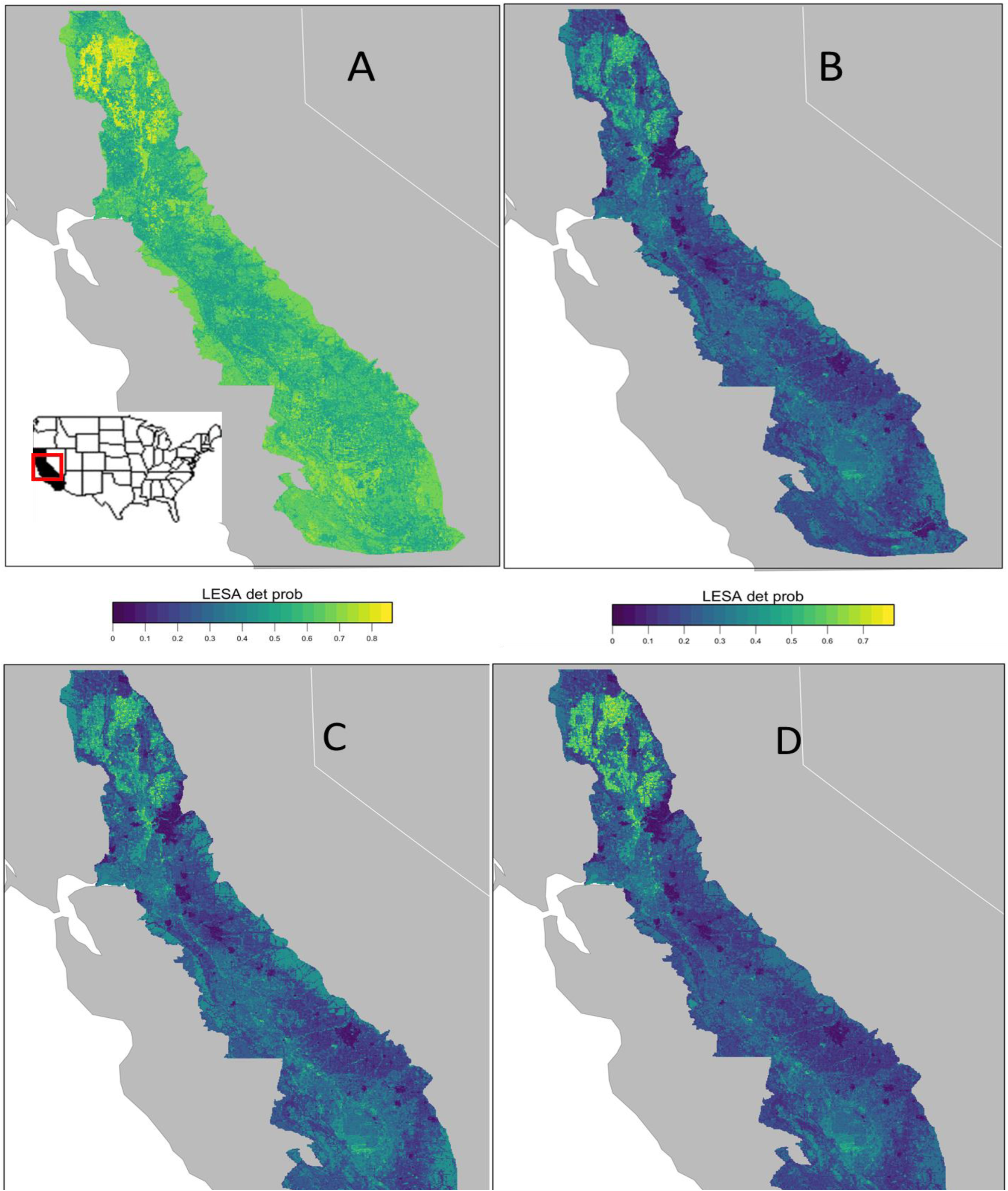

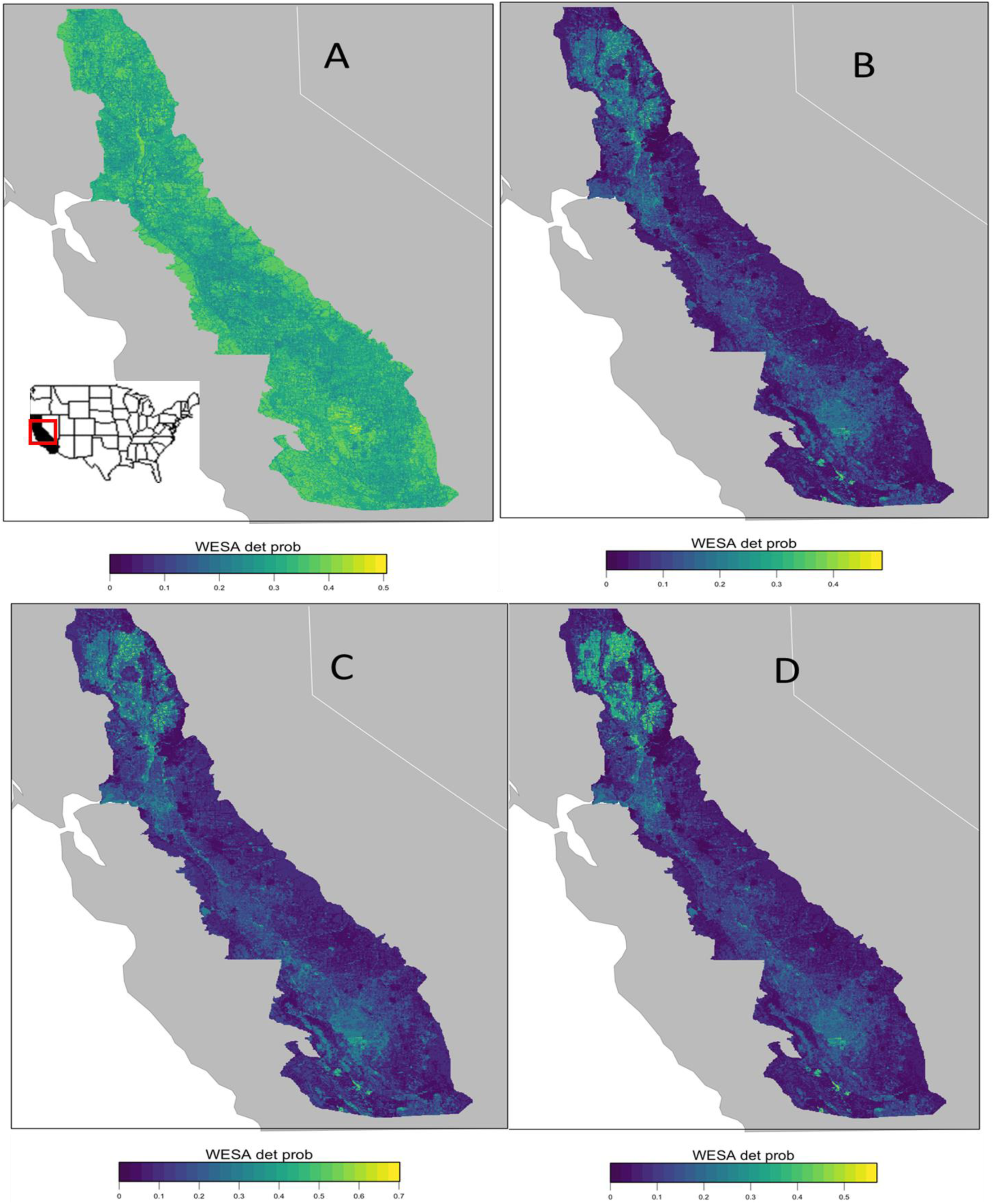

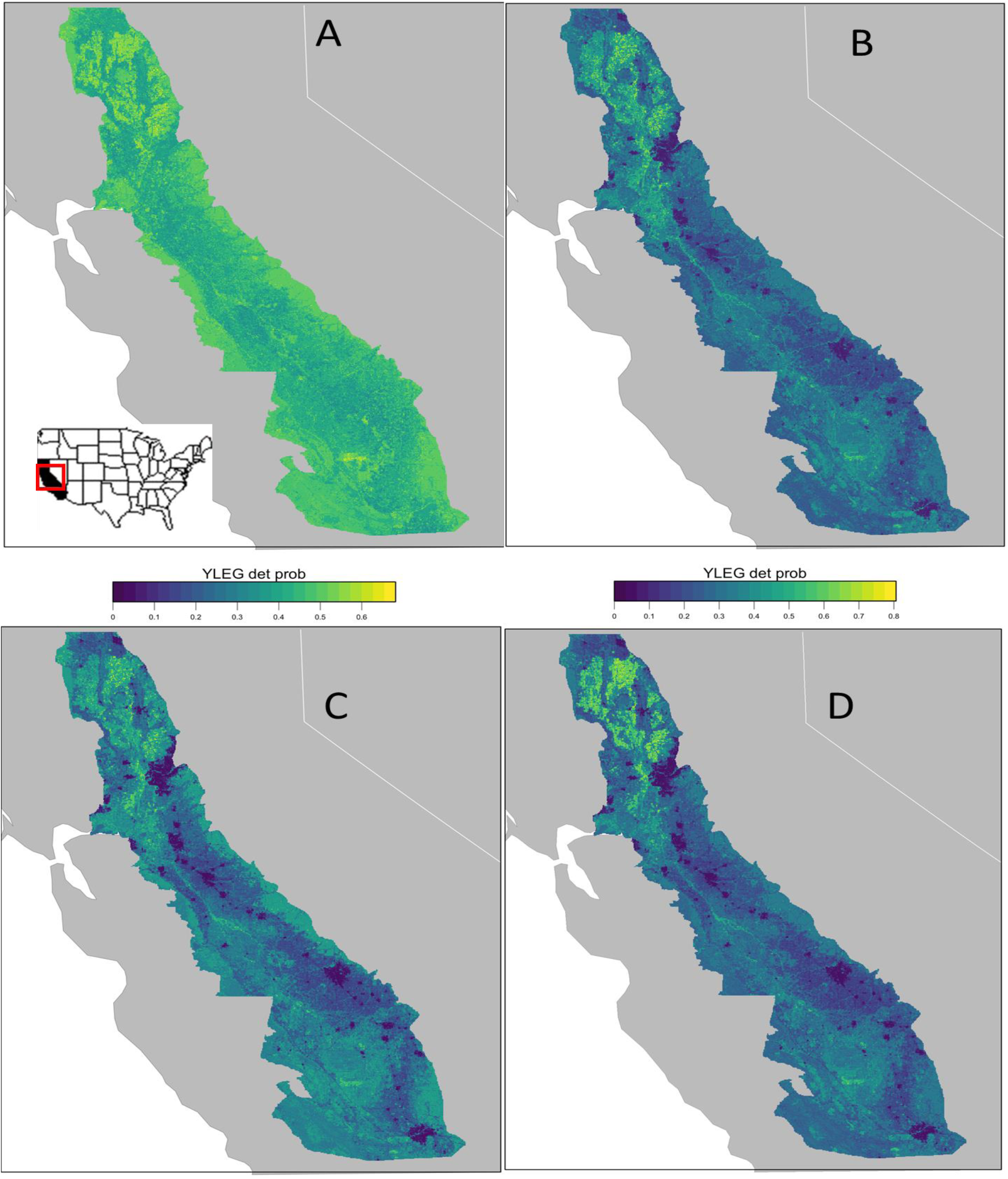

## Supplementary Information 4

Variable Importance for American Avocet, Dowithcer sp., Long-billed Curlew, Least Sandpiper, Western Sandpiper, and Yellowlegs sp. These figures for Dunlin appear in the main text.

Important variables selected by the model trained on each data set for American Avocet as selected by Gini importance index; TNC point counts alone (A), eBird checklists alone (B), and the combined dataset of TNC point counts and eBird checklists (C). Please note the different scales for each panel, and that this only represents the importance of each variable, not the direction of its relationship with probability of detection.

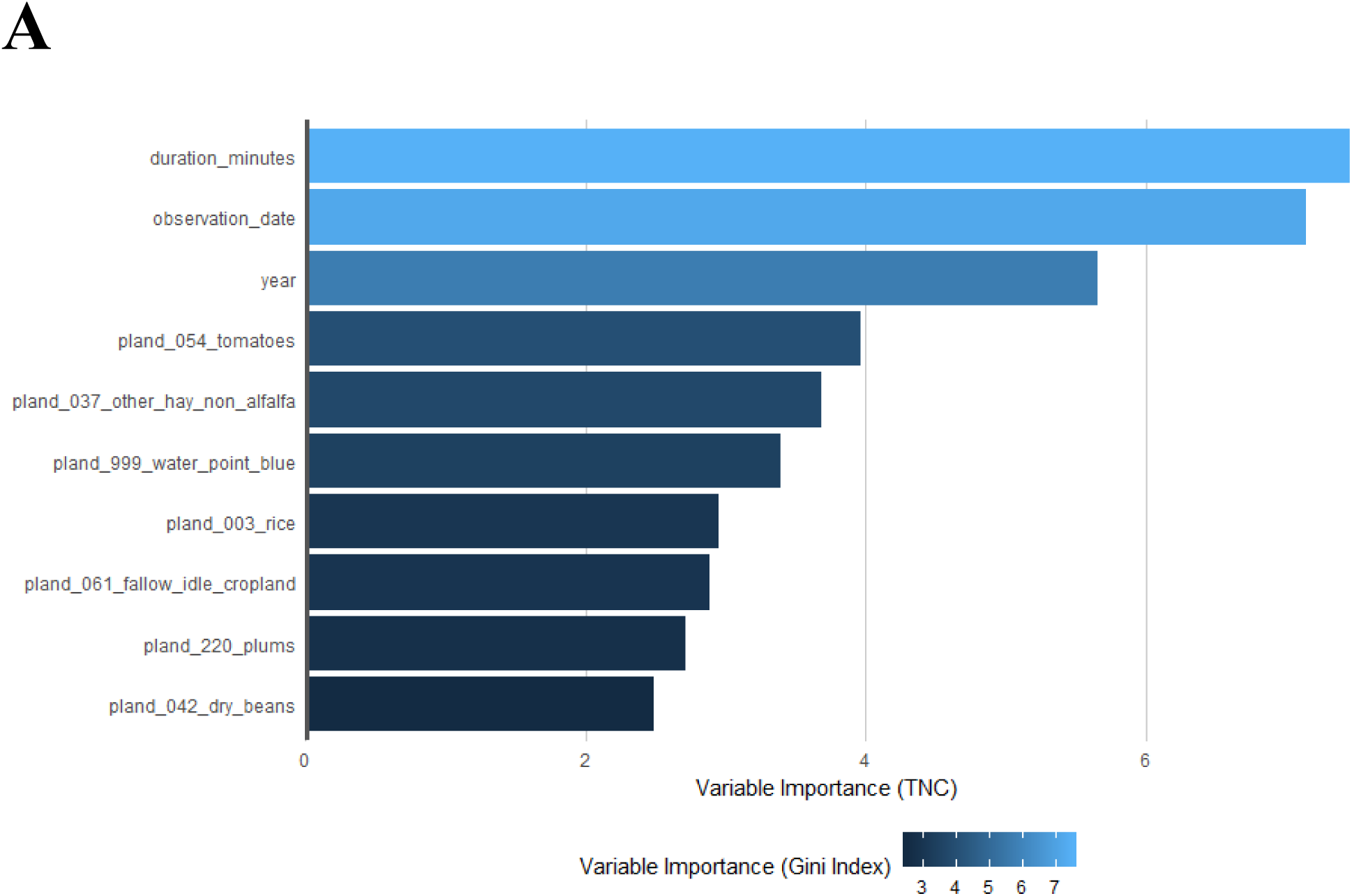

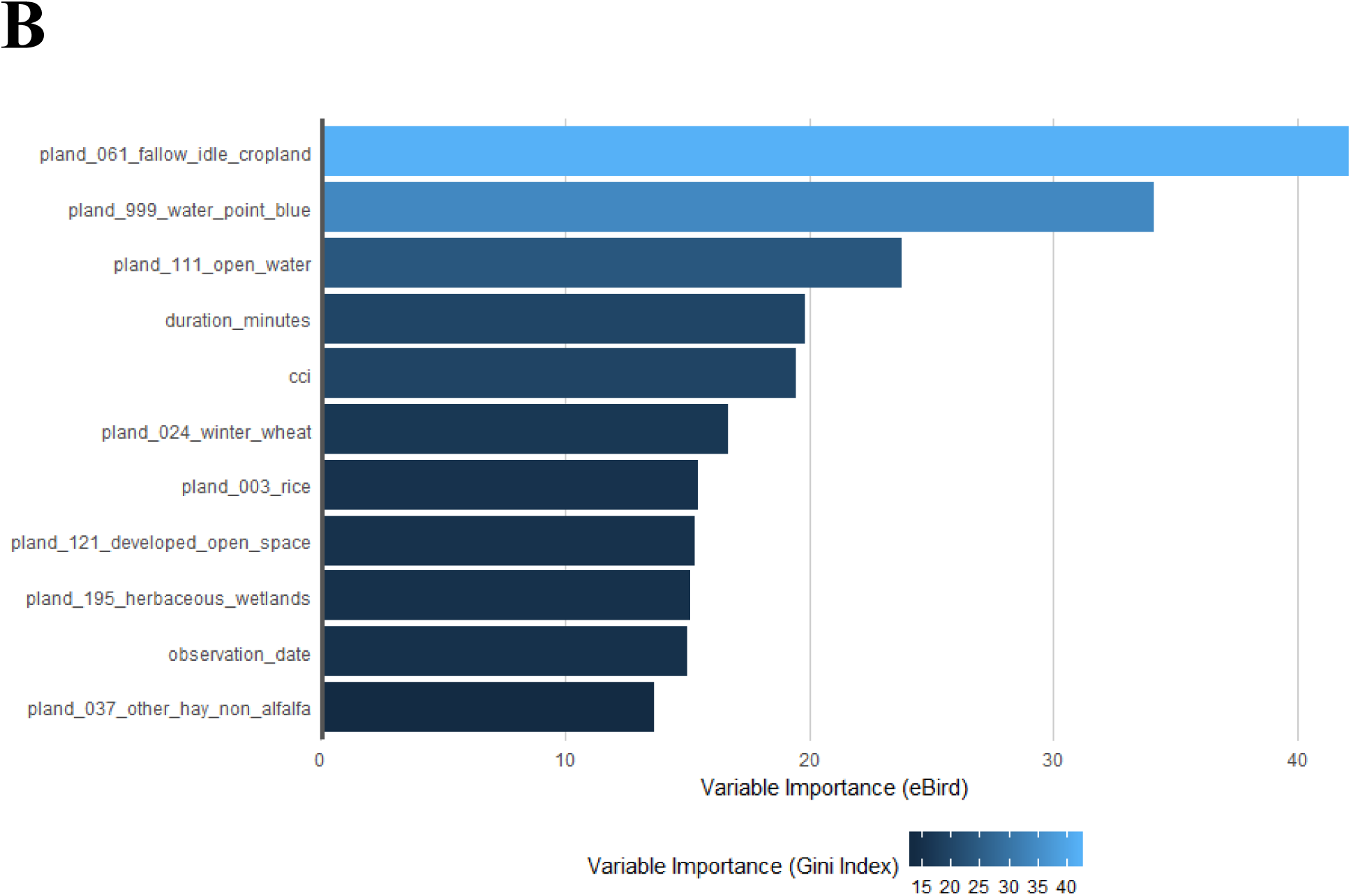

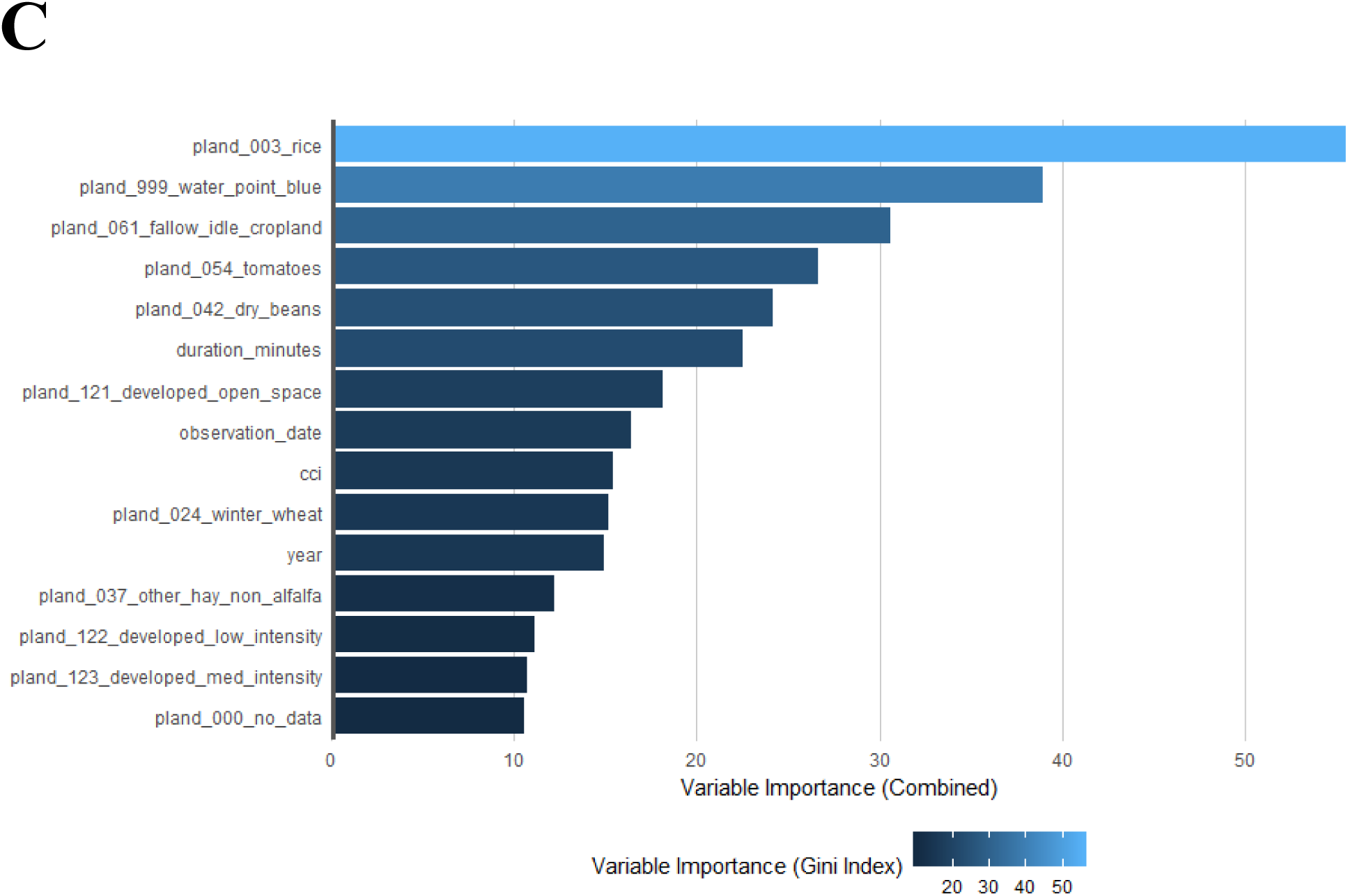

Important variables selected by the model trained on each data set for short- and long-billed dowitcher combined as selected by Gini importance index; TNC point counts alone (A), eBird checklists alone (B), and the combined dataset of TNC point counts and eBird checklists (C). Please note the different scales for each panel, and that this only represents the importance of each variable, not the direction of its relationship with probability of detection.

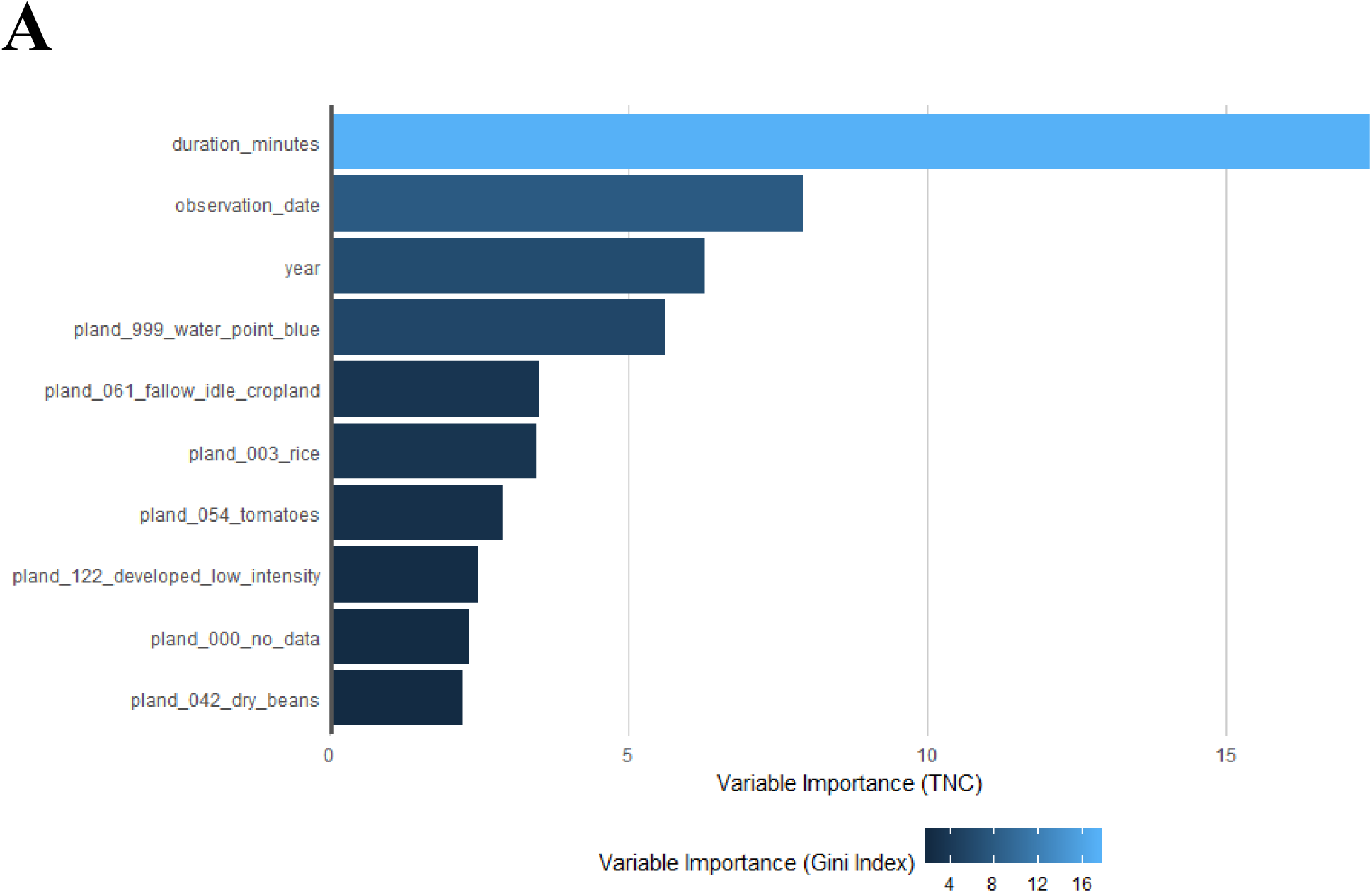

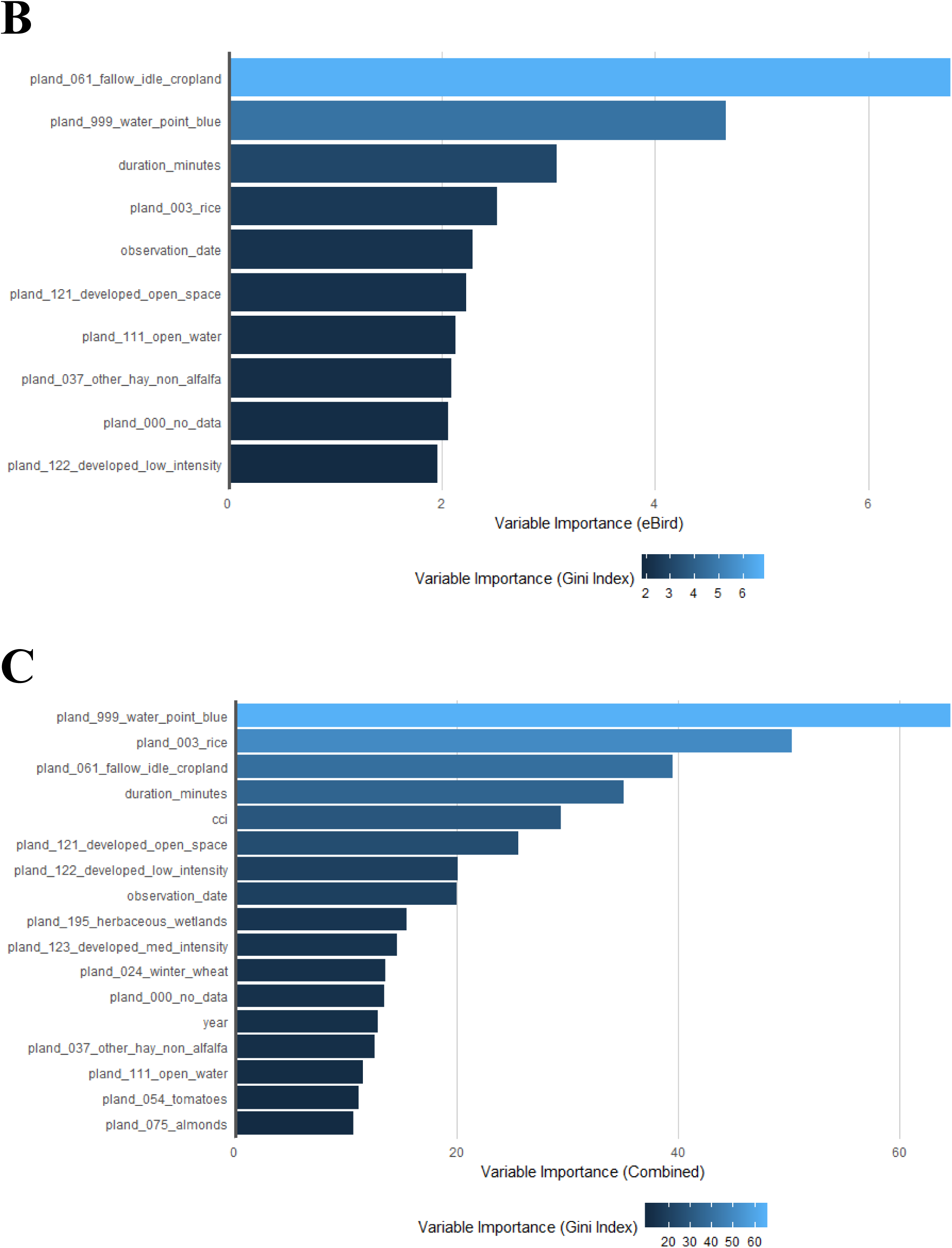

Important variables selected by the model trained on each data set for long-billed curlew as selected by Gini importance index; TNC point counts alone (A), eBird checklists alone (B), and the combined dataset of TNC point counts and eBird checklists (C). Please note the different scales for each panel, and that this only represents the importance of each variable, not the direction of its relationship with probability of detection.

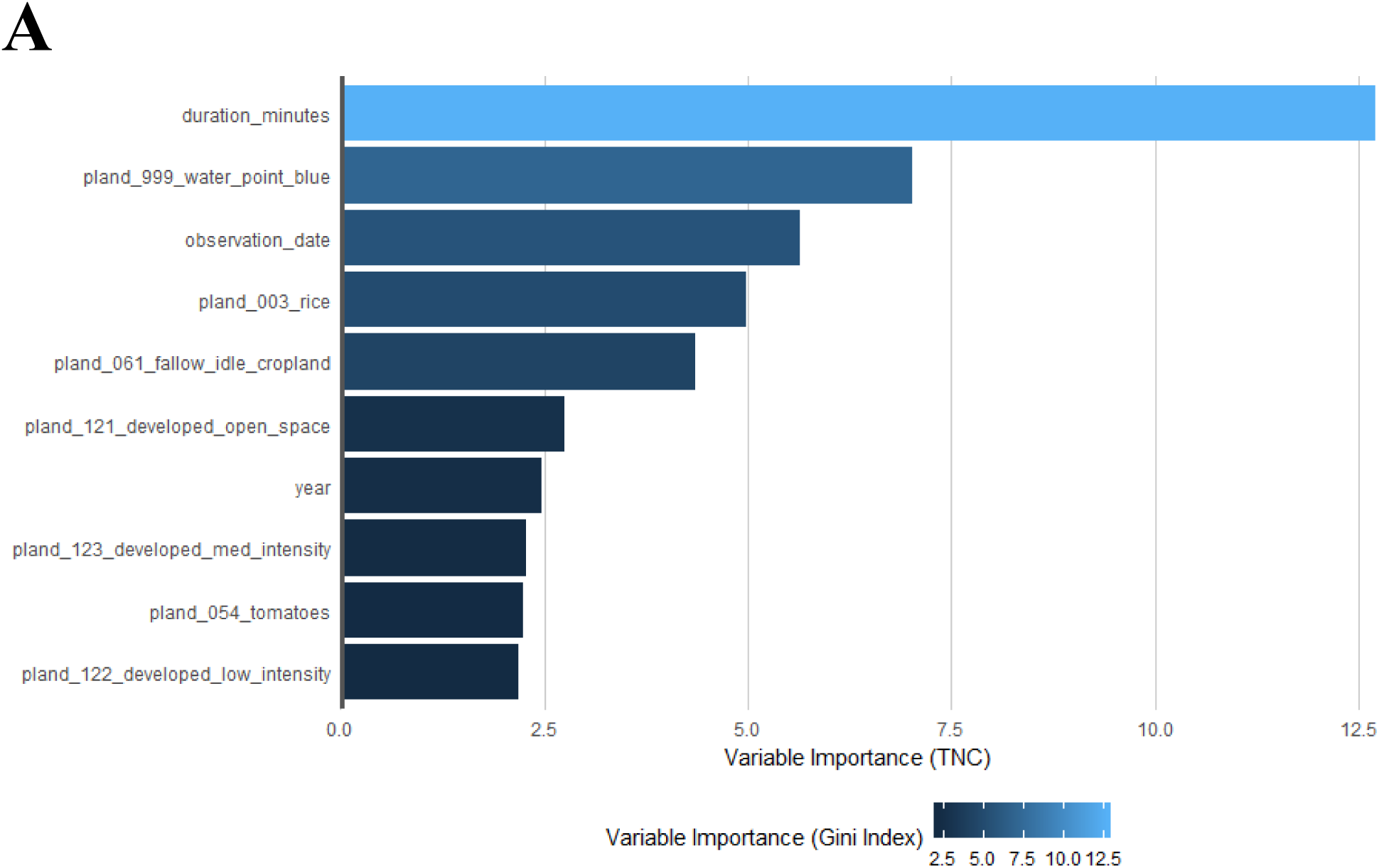

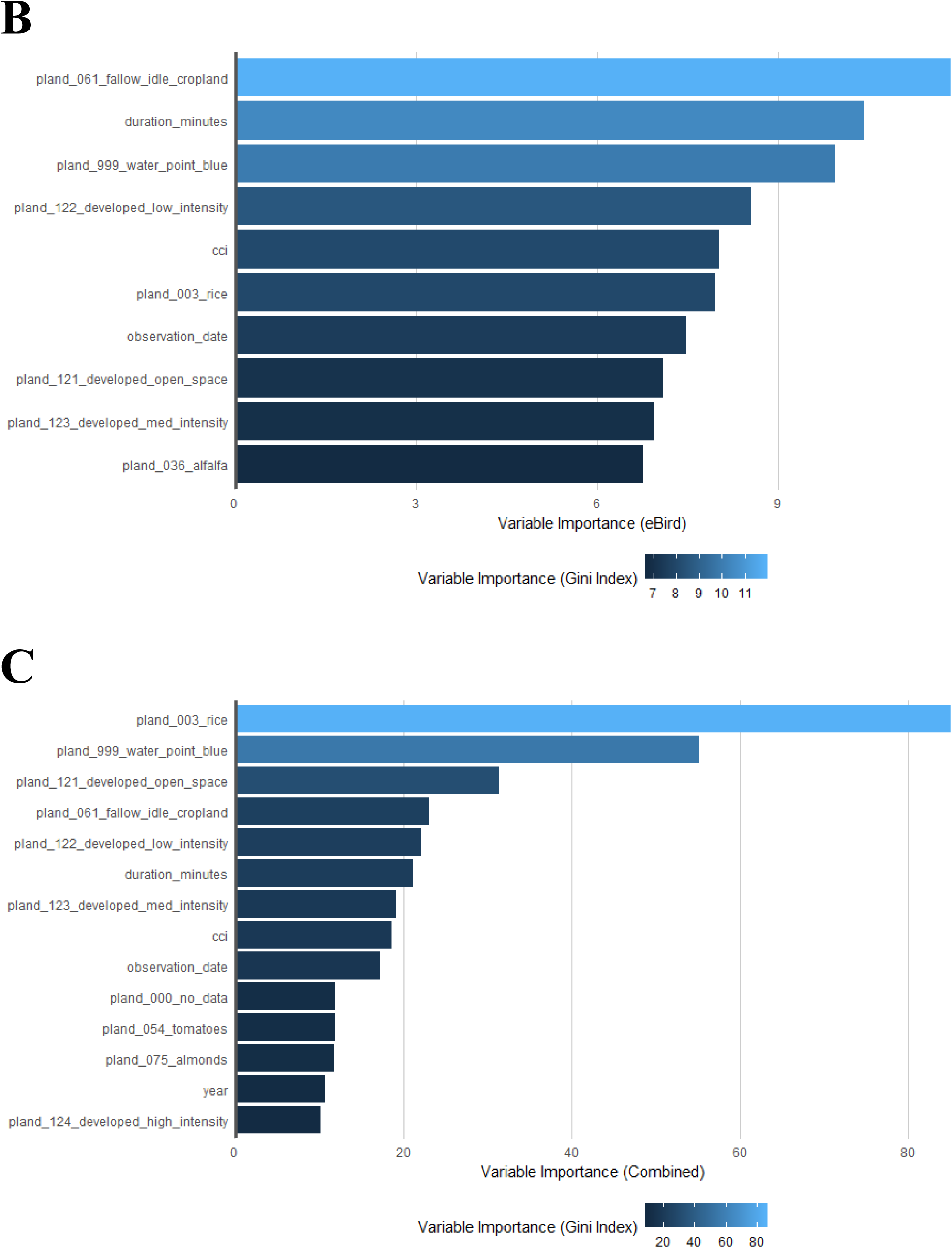

Important variables selected by the model trained on each data set for least sandpiper as selected by Gini importance index; TNC point counts alone (A), eBird checklists alone (B), and the combined dataset of TNC point counts and eBird checklists (C). Please note the different scales for each panel, and that this only represents the importance of each variable, not the direction of its relationship with probability of detection.

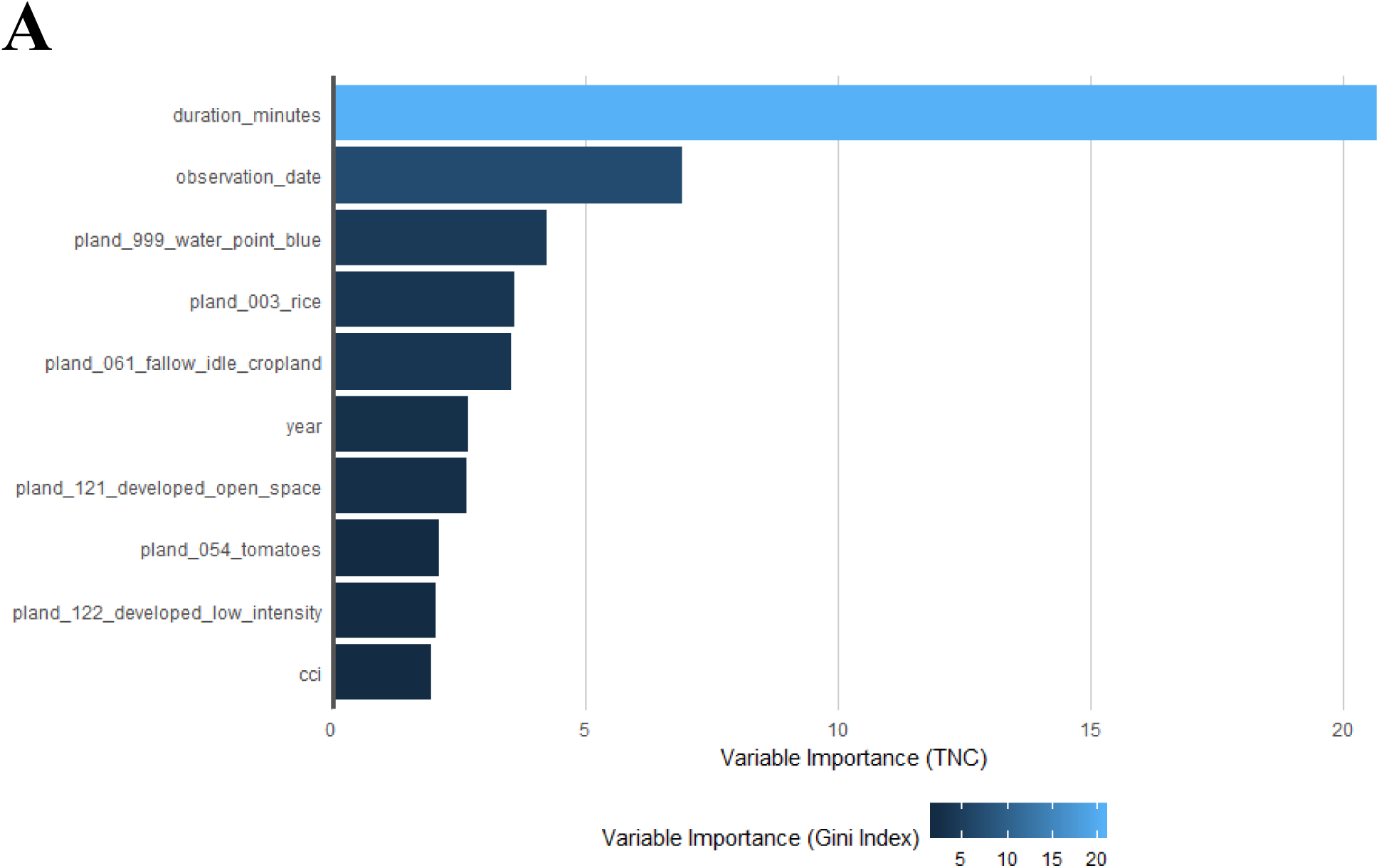

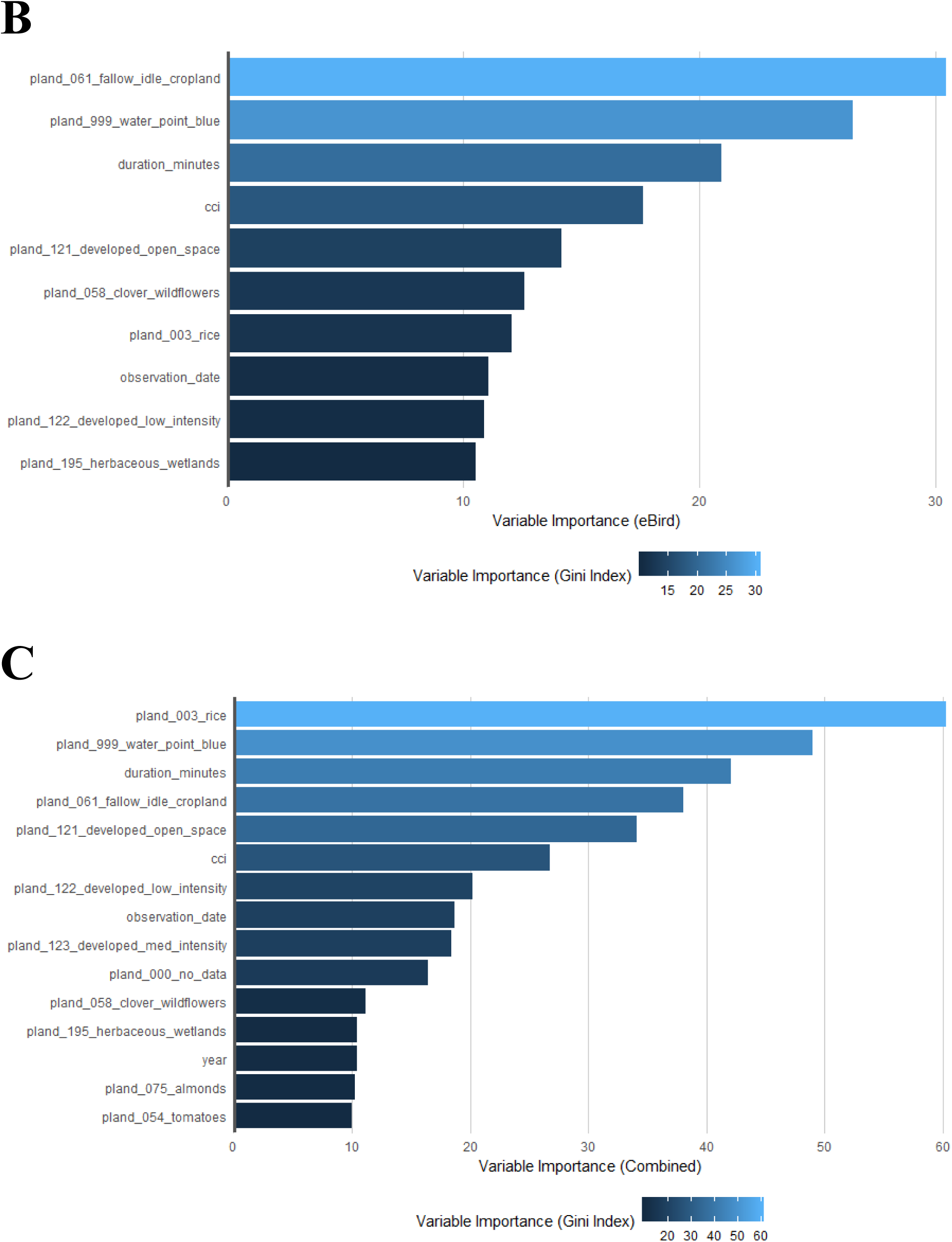

Important variables selected by the model trained on each data set for western sandpiper as selected by Gini importance index; TNC point counts alone (A), eBird checklists alone (B), and the combined dataset of TNC point counts and eBird checklists (C). Please note the different scales for each panel, and that this only represents the importance of each variable, not the direction of its relationship with probability of detection.

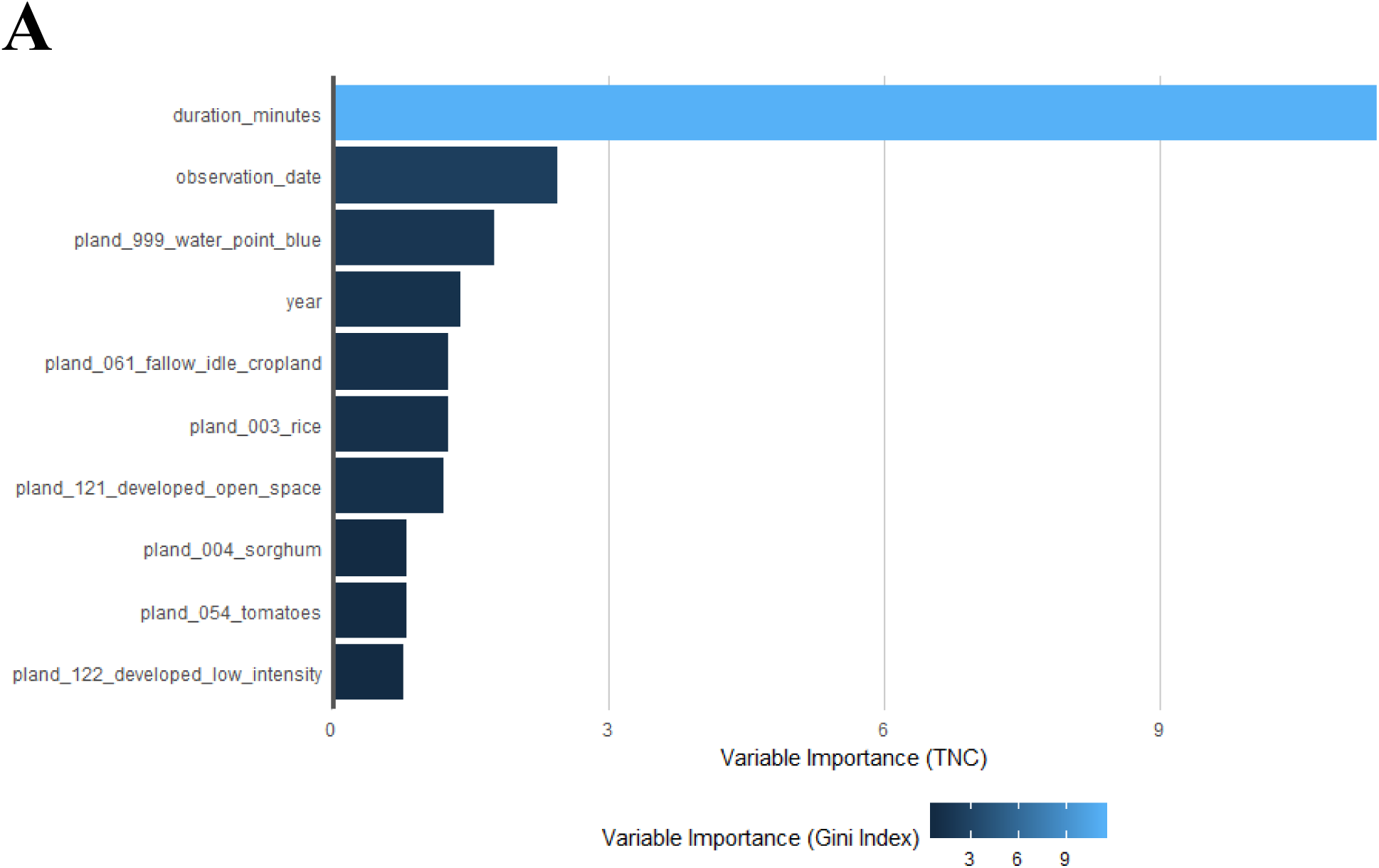

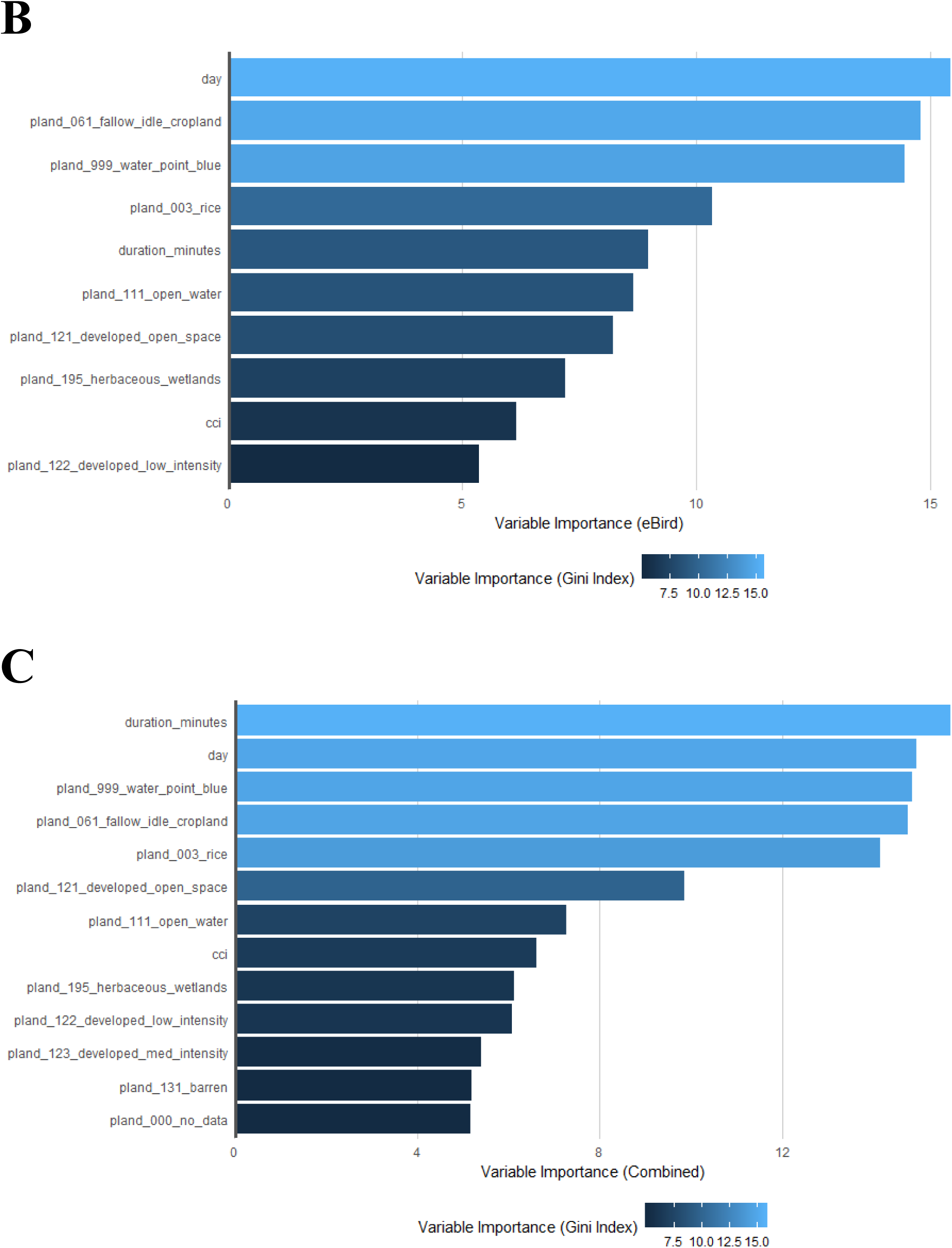

Important variables selected by the model trained on each data set for greater and lesser yellowlegs combined as selected by Gini importance index; TNC point counts alone (A), eBird checklists alone (B), and the combined dataset of TNC point counts and eBird checklists (C). Please note the different scales for each panel, and that this only represents the importance of each variable, not the direction of its relationship with probability of detection.

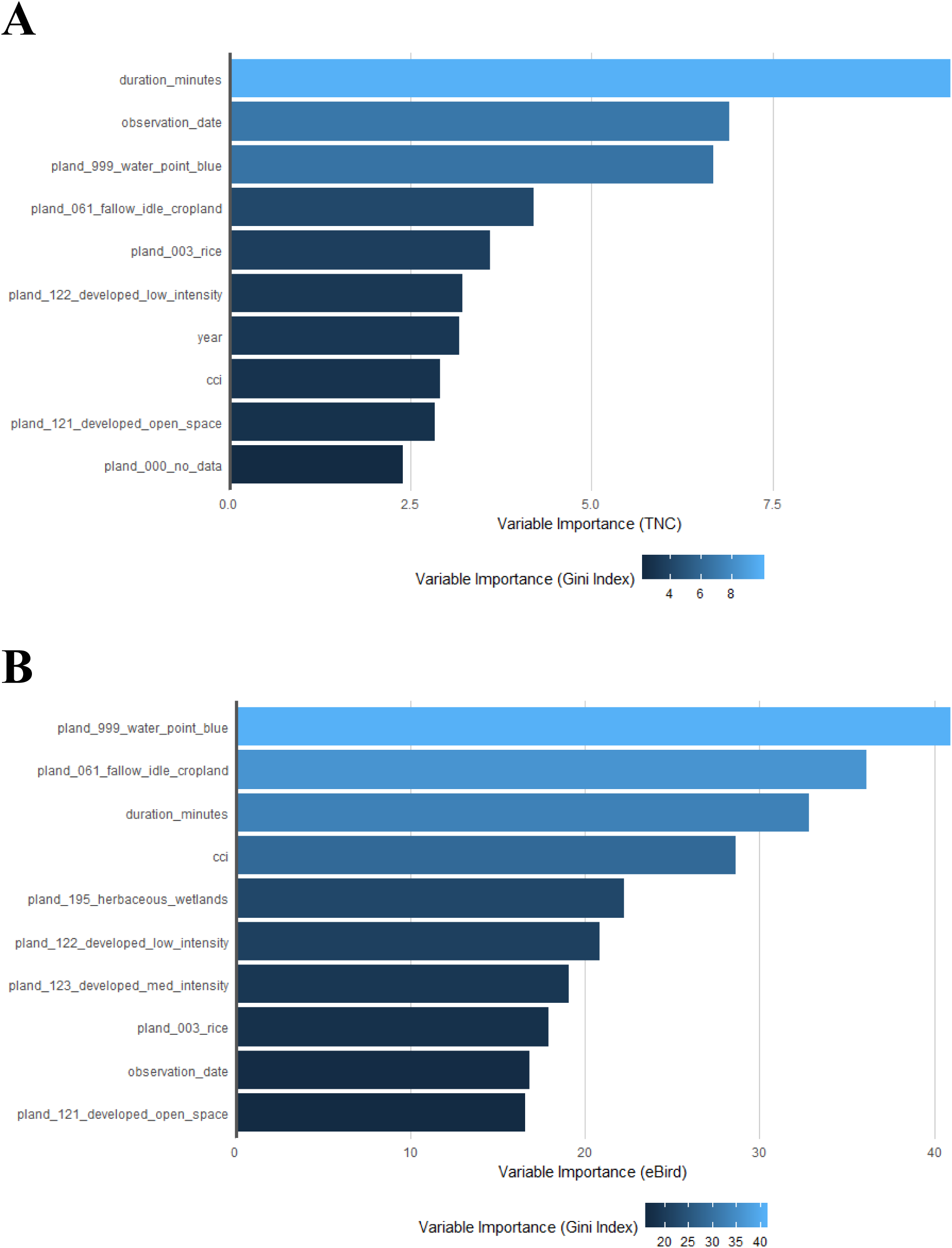

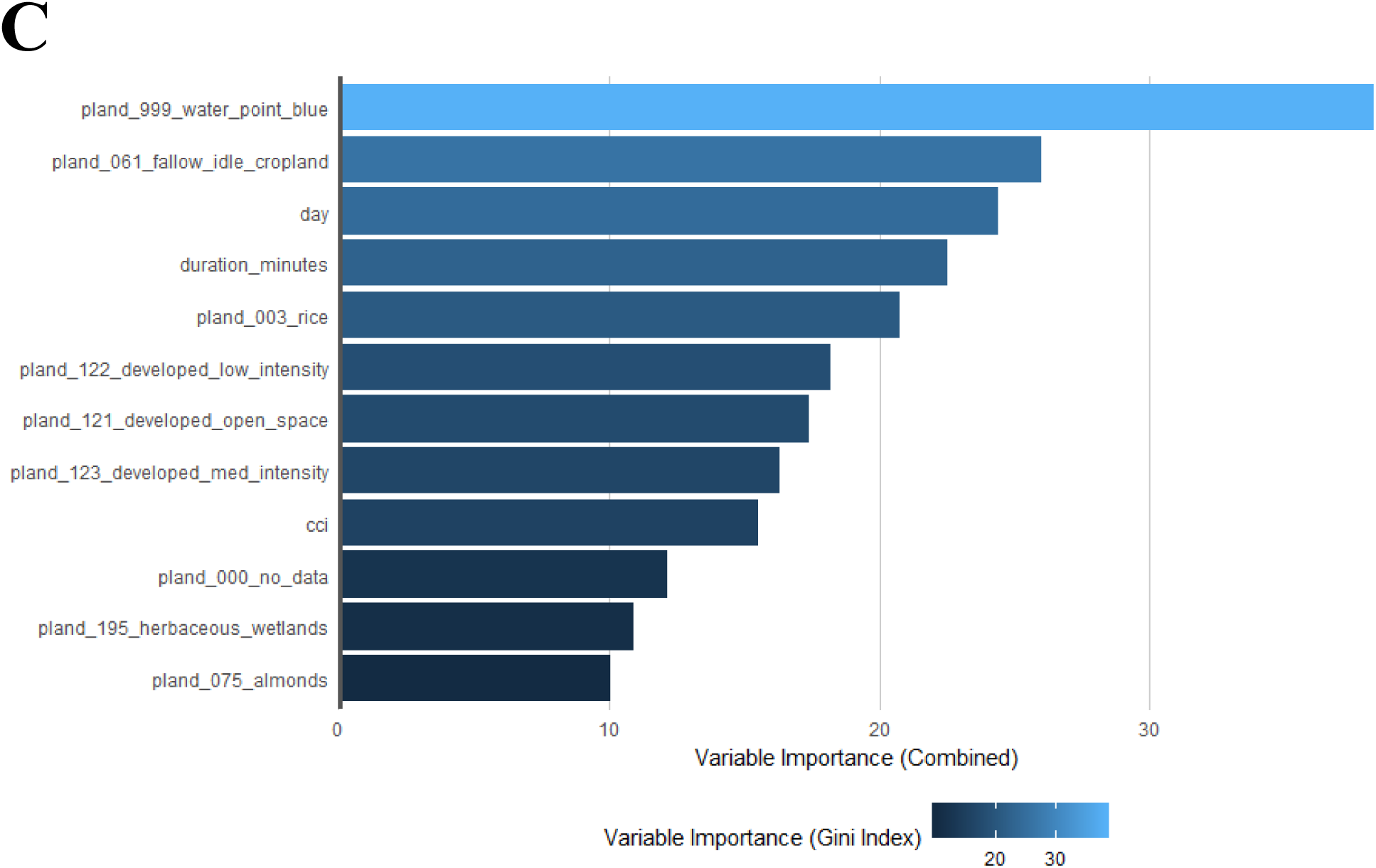

## Notes

#### Summary of Updates

Included title and abstract as submitted to journal for review. Added website for Water Tracker data. Added acknowledgements.

https://doi.org/10.5061/dryad.724vn

https://ebird.org/data/download

https://www.nass.usda.gov/Research_and_Science/Cropland/Release/index.php

https://data.pointblue.org/apps/autowater/

